# The genetic structure and adaptation of Andean highlanders and Amazonian dwellers is influenced by the interplay between geography and culture

**DOI:** 10.1101/2020.01.30.916270

**Authors:** Victor Borda, Isabela Alvim, Marla M Aquino, Carolina Silva, Giordano B Soares-Souza, Thiago P Leal, Marilia O Scliar, Roxana Zamudio, Camila Zolini, Carlos Padilla, Omar Cáceres, Kelly Levano, Cesar Sanchez, Omar Trujillo, Pedro O. Flores-Villanueva, Michael Dean, Silvia Fuselli, Moara Machado, Pedro E. Romero, Francesca Tassi, Meredith Yeager, Timothy D. O’Connor, Robert H Gilman, Eduardo Tarazona-Santos, Heinner Guio

## Abstract

Western South America was one of the worldwide cradles of civilization. The well known Inca Empire was the *tip of the iceberg* of a cultural and biological evolutionary process that started 14-11 thousand years ago. Genetic data from 18 Peruvian populations reveal that: (1) The between-population homogenization of the central-southern Andes and its differentiation with respect to Amazonian populations of similar latitudes do not extend northward. Instead, longitudinal gene flow between the northern coast of Peru, Andes and Amazonia accompanied cultural and socioeconomic interactions revealed by archeological studies. This pattern recapitulates the environmental and cultural differentiation between the fertile north, where altitudes are lower; and the arid south, where the Andes are higher, acting as a genetic barrier between the sharply different environments of the Andes and Amazonia (2). The genetic homogenization between the populations of the arid Andes is not only due to migration during the Inca Empire or the subsequent colonial period. It started at least during the earlier expansion of the pre-Inca Wari Empire (600-1000 YBP) (3) This demographic history allowed for cases of positive natural selection in the high and arid Andes vs. the low Amazon tropical forest: in the Andes, *HAND2-AS1* (heart and neural crest derivatives expressed 2 antisense RNA1, related with cardiovascular function) and *DUOX2* (dual oxidase 2, related to thyroid function and innate immunity) genes; in the Amazon, the gene encoding for the CD45 protein, essential for antigen recognition by T/B lymphocytes in viral-host interaction, consistent with the *host-virus arms race* hypothesis.

## Main text

Western South America was one of the cradles of civilization in the Americas and the world ^1^. When the Spaniard conqueror Francisco Pizarro arrived in 1532, the pan-Andean Inca Empire ruled in the Andean region and had achieved levels of socioeconomic development and population density unmatched in other parts of South America. However, the Inca Empire, which lasted for around 200 years, with its emblematic architecture such as Machu Picchu and the city of Cuzco, was just the *tip of the iceberg* of a millenary cultural and biological evolutionary process ^2,3^. This process started with the peopling of the region (hereafter called *western South America*), that occurred 14–11 thousand years ago ^4–6^, involving the entire Andean region and its adjacent and narrow Pacific Coast.

Tarazona-Santos et al. ^7^ proposed that cultural exchanges and gene flow along time have led to a relative genetic, cultural, and linguistic homogeneity between the populations of western South America compared with those of eastern South America (a term that hereafter refers to the region adjacent to the eastern slope of the Andes and eastward, including the Amazonia), where populations remained more isolated from each other. For instance, only two languages (Quechua and Aymara) of the Quechumaram linguistic stock predominate in the entire Andean region, whereas in eastern South America natives speak a different and broader spectrum of languages classified into at least four linguistic families ^3,7,8^. This spatial pattern of genetic diversity and its correlation with geography, environmental, linguistic and cultural diversity was confirmed, enriched and rediscussed by us and others ^2,3,7–13^.

There are pending issues: First, whether the dichotomic organization of genetic variation characterized by the between-population homogeneous Southern Andes vs. between-population heterogeneous Central Amazon, extends northward. This is important because scholars from different disciplines emphasize that western South America is not latitudinally homogeneous, differentiating a northern and in general lower and wetter fertile Andes and a southern, higher and more arid Andes ^14^. These environmental and latitudinal differences are correlated with demography and culture, including different spectra of domesticated plants and animals. Indeed, the development of agriculture, of the first urban centers such as Caral ^1^ and its associated demographic growth, occurred earlier in the northern Fertile Andes (around 5ky ago) than in the southern arid Andes (and their associated Coast), with products such as cotton, beans, and corn domesticated in the fertile north and the potato and South American camelids in the arid south ^14^. In human population genetics studies, the region where the between-population homogeneity was ascertained by Tarazona-Santos et al.^7^ was the arid Andes. Consequently, here we test **(i) whether the between-population homogenization of Western South America, and the dichotomy Arid Andes/Amazonia extends to the northward Fertile Andes associated regions?** To address this and the below questions, we used data from Harris et al.^3^ for 74 indigenous individuals and an additional 289 unpublished individuals from 18 Peruvian populations, genotyped for ~2.5 million SNPs (Figure 1 and Table S1). We created three datasets with different SNP densities and populations, including data from ^15–18^ (Figure S1, Tables S2 and S3 and Supporting information-SI). Institutional Review Boards of participants institutions approved this research.

**Figure 1.**
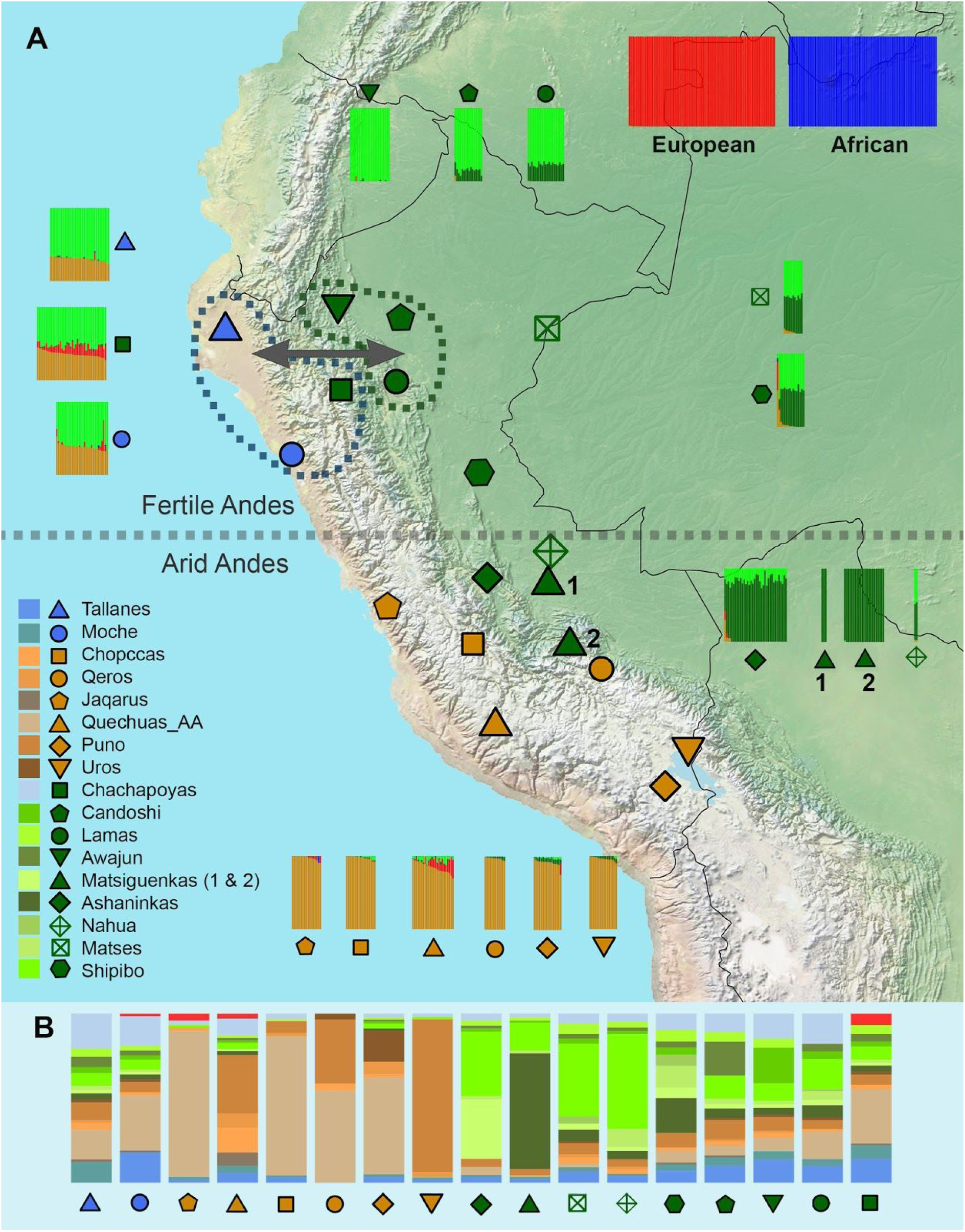
Genetic structure and gene flow of Western South American natives. Eighteen native groups along the Coast, Andes, and Amazon regions were sampled. A) Grey dashed line in the center shows the division between Fertile Andes and Arid Andes ^14^. We showed the geographical distribution coupled with ADMIXTURE patterns for the lowest cross-validation value (K=5) using the dataset of 1.9M unlinked SNPs (including Native Americans, Europeans, and Africans). Blue and green dashed lines delimited the groups that showed highly significant value for the gene flow test (|Z score| > 4). Matsiguenka 1= Matsiguenkas-Sepahua, Matsiguenkas 2= Matsiguenkas-Shimaa B) Haplotype based inference of ancestry profile for each Native American population, each bar corresponds to the ancestry composition for a native population. For this analysis, Matsiguenka samples were merged into one. Colors for the ancestry profile correspond to the proportion of DNA shared between the population and donor groups detailed in the legend of section A.

By applying ADMIXTURE ^19^ (Figures 1, S2, S4, S6) and principal component analyses (Figures S3, S5, S7), as well as haplotype-based methods ^20,21^ (Figures S8-S13), we confirmed that populations in the Arid Andes are genetically homogeneous, appearing as an almost panmictic unit, and with an ancestry pattern differentiated with respect to Amazonian populations of similar latitudes (Figure 1). Conversely, populations of the northern Coast (Moches), in the north Coast/Andes interface (Tallanes) and in the northern Amazon Yunga (the rainforest transitional region between the Andes and the Lower Amazonia) (Chachapoyas), share the same ancestry profile between them (Figures 1, S8-S13), which is different from the populations from the Arid Andes. Thus, the between-population homogenization of the Arid Andes and its differentiation respect to Amazonian populations of similar latitudes do not extend northward and is not characteristic of all Western South America. Instead, the genetic structure of Western South Amerindian populations recapitulates the environmental and cultural differentiation between northern Fertile Andes and the southern Arid Andes.

A second open issue is the evolutionary relationship between Andean and Amazonian populations, particularly with the culturally, linguistically, and environmentally different neighboring populations of the Amazon Yunga. Harris et al. ^3^ inferred that Andean and Amazonian populations diverged around 12,000 years ago. Archaeological findings of recent decades have rejected the traditional view of the Amazonian environment as incompatible with complex pre-Columbian societies, and have revealed that the Amazonian basin has produced the earliest ceramics of South America, that endogenous agricultural complex societies have developed there, and that population sizes were larger than previously thought ^22^. Population genetics studies by Barbieri et al. ^23^ have reported episodes of gene flow in the Amazonia, which suggest that Amazonian populations were not necessarily isolated groups. Moreover, people living in the Peruvian Coast, the Andes, and the Amazon Yunga had cultural and commercial interactions during the last millennia, sharing practices such as sweet potato and manioc cultivation, ceramic iconography and styles (e.g. Tutishcanyo, Kotosh, Valdivia and Corrugate) and traditional coca chewing ^24^. Therefore, here we address (ii) **whether gene flow accompanied the cultural and socioeconomic interactions between Andean and Amazon Yunga populations**?

Haplotype-based inferences ^20,21^ (Figures 1 and S11-S13) and statistical tests of treeness ^25^ (Figures 1 and S14-S17) show genetic signatures of gene flow between Coastal/Andean and Amazon Yunga populations in latitudes of the northern fertile Andes, but not in the southern arid Andes. Thus, longitudinal gene flow between the North Coast, Andes, and Amazonia accompanied the well documented cultural and socioeconomic interactions, recapitulating the differentiation between the fertile north, where altitudes are lower; and the arid south, where the Andes altitudes are higher and may have acted as a barrier to gene flow, imposing a sharper environmental differentiation between the Andes and the Amazon Yunga. Formal tests show that in latitudes of the fertile north, gene flow includes important ethnic groups such as the current Chachapoyas of the Amazon Yunga, as well as eastward Lower Amazonian populations such as of the Jivaro linguistic family (Awajun and Candoshi) and Lamas (Figure 1 and S18-S21).

Despite some controversy about definitions and chronology, archeologists identify a unique cultural process in Western South America, which include three temporal *Horizons*: Early, Middle, and Late, that corresponds to periods of cultural dispersion involving a wide geographic area ^26^ (Figure 2). In particular, the Middle and Late Horizons are associated with the expansions of the Wari (~1400 to 1000 YBP) and Inca (~524 to 466 YBP) states, respectively ^27–29^. Isbell ^28^ has suggested that the Wari expansion has been associated with the spread of the Quechua language in the Central Andes and the Wari were pioneers in developing a road system in the Andes, called *Wari ñam*, which was later used as a base by the Incas to develop their network roads (the *Qapaq ñam*) (Lumbreras et al. 2015). The between-population homogeneity currently observed in the arid Andes implies high levels of gene flow in this region, which are commonly associated with the Inca Empire ^26^. A relevant question is: **(iii) when this between-population genetic homogenization started in the context of the arid Andean chronology.** Particularly, is this is a phenomenon restricted to the time of the Inca Empire or did it extend backward to the Middle/Wari Horizon?

**Figure 2.**
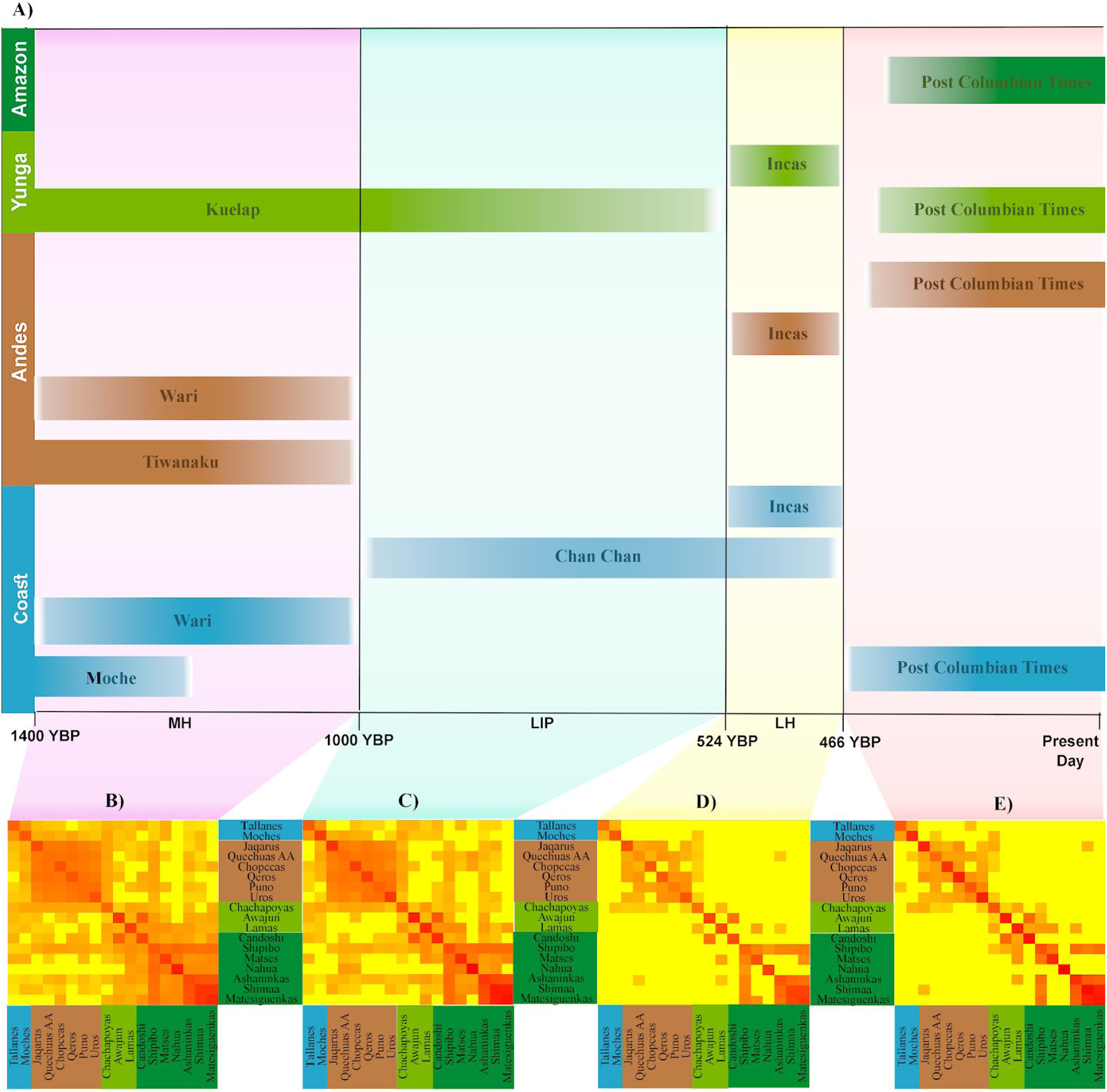
Evolution of IBD sharing between the Pacific Coast, Central Andes, Amazon Yunga and Amazon and its relationship with archaeological chronology of the Andes. A) Figure 2 of Scliar et al.^2^ adapted, showing key historical events (cultures and archeological sites) of Peruvian history in each region. YBP: Years Before Present, LH: Late Horizon, LIP: Late Intermediate Period, MH: Middle Horizon. B), C) and D) are heat-maps of the average pairwise relatedness (Baharian et al 2016) among Native Americans of the Natives 1.9M dataset. Each heatmap represents an interval of IBD segment lengths, which correspond to interval times (Palamara et al. 2012). B) Interval from 3.2-4.2 cM corresponding to 50-36 generations ago. C)Interval from 4.2-7.8 cM corresponding to 36 to 19 generations ago. D) Interval 7.8-9.3 cM corresponding to 19-16 generations ago. E) The last interval for all segments longer than 9.3 cM corresponds to 16-0 generations ago.

We analyzed the distribution of IBD-segment lengths between individuals of different arid Andean populations, which is informative for the dynamics of past gene flow ^3,30^, and observed a signature of gene flow in the interval 1400 to 1000 YBP, that is within the Wari expansion in the Middle Horizon (Figure 2). Thus, the homogenization of the Central Arid Andes is not only due to migrations during the Inca Empire or later during the Spanish Viceroyalty of Peru, when migrations (often forced) occurred ^31^. The Wari expansion (1400 to 1000 YBP) was also accompanied by intensive gene flow whose signature is still present in the between-population genetic homogeneity of the arid central Andes region. Because IBD analysis on current individuals does not allow for inferences of gene flow that occurred more than 75 generations ago ^30^, ancient DNA analysis at the population level will be necessary to infer if the between-population homogenization of the Andes started even earlier.

Native Americans had to adapt to different and contrasting environments and stress. The high and arid Andes is characterized by high UV radiation, cold, dryness, and hypoxia (a stress that does not allow for cultural adaptations and requires biological changes) ^32,33^. The Amazon has a low incidence of light, a warm and humid climate typical of the rainforest and high biodiversity, including human pathogens ^34^. Populations from the high and arid Andes and from the Amazon (Figure 1) settled in these contrasting environments more than 5000 years ago ^35^ and show little evidence of gene flow between them (i.e. that would homogenize allele frequencies, potentially concealing the effect of diversifying natural selection). Thus, we performed genome-wide scans in these two groups of populations using two tests of positive natural selection: (i) Population Branch Statistics (PBSn) comparing arid Andeans (Chopccas, Quechuas, Qeros, Puno, Jaqarus, and Uros) vs. Amazonian populations (Ashaninkas, Matsiguenkas, Matses and Nahua) with CDX population (Chinese Dai in Xishuangbanna, China) from 1000 Genomes as an outgroup ^36^ and (ii) long-range haplotypes (xpEHH)^37^ estimated with the same two groups of populations (Supplementary text, Section 4, Figure 3, S25-S27). The complete lists of SNPs with high PBSn and xpEHH statistics for Andean and Amazonian populations are in Tables S4-S7.

**Figure 3.**
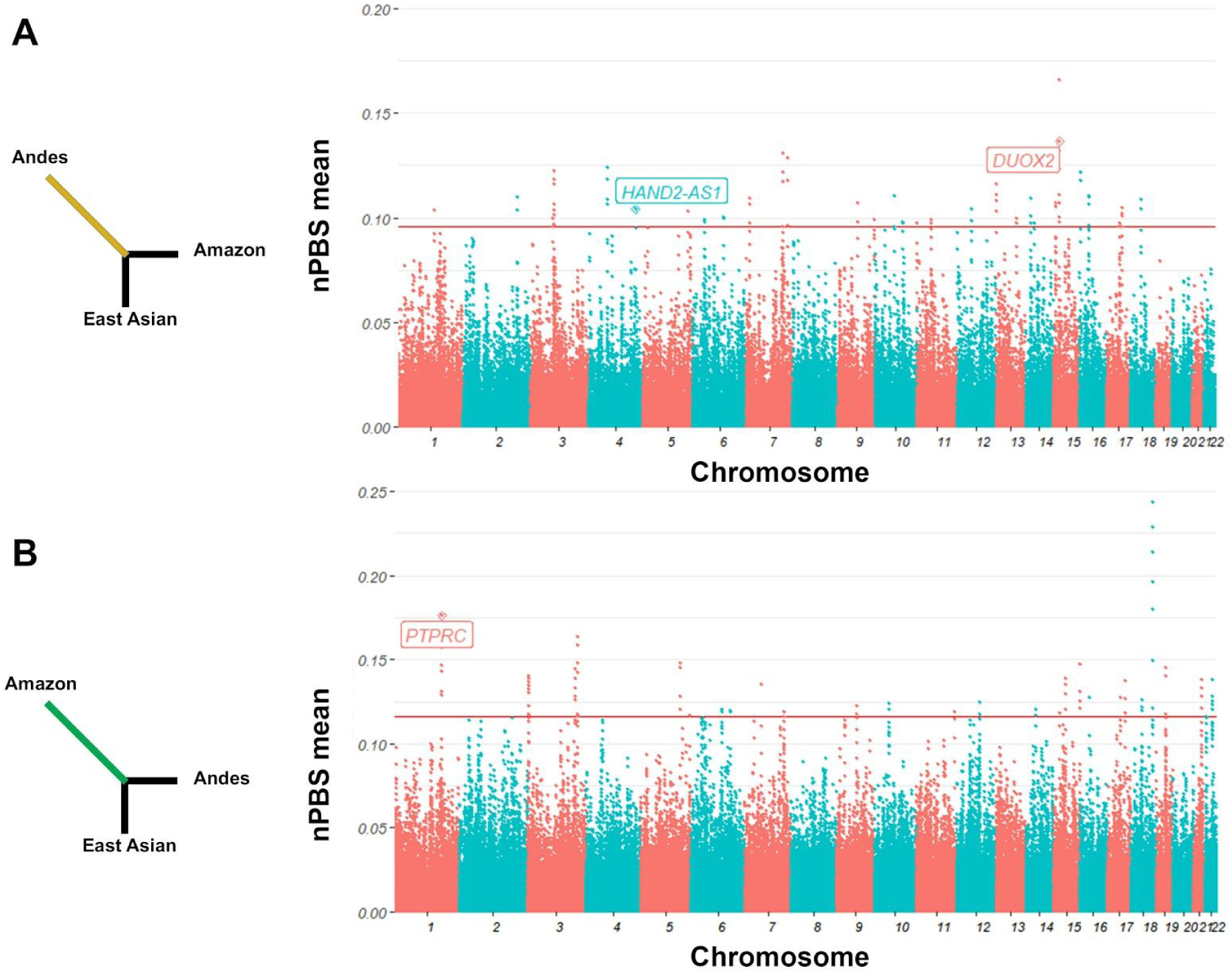
Manhattan plot of the Population Branch Specific (PBSn) estimated as a sliding windows of 20 SNPs with 5 SNPs of overlap in Andean (A) and Amazon (B) populations. We tagged PBSn hits that correspond to genes and show a significant xpEHH statistics. The horizontal red line shows 95.95th percentile of PBSn values.

The gene with the consensually strongest signal of adaptation (both from PBSn and xpEHH statistics) to the Andean environment (Figure 3, Table S4) is the long non-coding RNA gene *HAND2-AS1* (heart and neural crest derivatives expressed 2 RNA antisense 1, chromosome 4), that modulates cardiogenesis by regulating the expression of the *HAND2* gene ^38,39^. HAND2-AS1 is located in the antisense 5’ region of HAND2 and contains 2 enhancers for this gene. A natural selection genome-wide scan ^36^ identified three genes related to the cardiovascular system in Andeans, including TBX5, which works together with HAND2 in reprogramming fibroblasts to cardiac-like myocytes ^40,41^. This information suggests (but does not demonstrate) that HAND2-AS1 signature of natural selection is related with cardiovascular cardiovascular adaptations. Andeans have cardiovascular adaptations to high altitude that differ from those of lowlanders exposed to hypoxia and from other highlanders, showing higher pulmonary vasoconstrictor response to hypoxia and lower resting middle cerebral flow velocity than Tibetans, and higher uterine artery blood flow than Europeans raised in high altitude and than lowlanders ^42^.

*DUOX2* (dual oxidase 2, chromosome 15) is the gene with the highest signal of adaptation to the Andean environment by PBSn analysis (Figure 3). It was reported as a natural selection target in the Andes by ^43,44^. DUOX2 encodes a trans-membrane component of an NADPH oxidase, which produces hydrogen peroxide (H_2_O_2_), and is essential for the synthesis of the thyroid hormone and for the production of microbicidal hypothiocyanite anion (OSCN–) during mucosal innate immunity response against bacterial and viral infections in the airways and intestines ^45,46^. Mutations in *DUOX2* produce inherited hypothyroidism ^47^. Here we report for the first time that: (i) The PBS signal for *DUOX2* comprises several SNPs, including 2 missense mutations (rs269868: C>T:Ser1067Leu, C allele frequencies: Amazon: 0.01, Andean: 0.53; rs57659670: T>C:His678Arg, C allele frequencies: Amazon: 0.01, Andean: 0.53), thus the ancestral allele has been positively selected in the Andes; (ii) bioinformatics analysis reveals that rs269868 is located in an A-loop aa1064-1078, which is a region of interaction of DUOX2 with its coactivator *DUOXA2*. Mutations in this region of the protein can affect the stability and maturation of the dimer and, consequently, the conversion of the intermediate product O_2_^□^ to the final product H_2_O_2_ and their released proportions ^48^. It is not clear if the *DUOX2* natural selection signal is related to thyroid function or innate immunity. Before the introduction of the public health program of supplementing manufactured salt with iodine, one of the environmental stresses of the Andes for human populations was iodine deficiency, which impairs thyroid hormone synthesis, increasing the risk of developing hypothyroidism, goiter, obstetric complications and cognitive impairment ^49,50^.

Natural selection studies in Amazon populations are scarce. Studies targeting rainforest populations in Africa and Asia have found natural selection signals in genes related to height and immune response ^51^. In the Amazon region, the strongest natural selection PBSn signal is in a long non-coding RNA gene on chromosome 18 with unknown function (Table S5). The second highest signal (that also shows a significant long-range haplotype signal) comes from the region around the gene *PTPRC*, which encodes the protein CD45, essential in antigen recognition by T and B lymphocytes, particularly in pathogen-host interaction, in particular for virus such as Human adenovirus type 19 ^52^, HIV-1-induced cell apoptosis ^53,54^ and susceptibility to Hepatitis C ^55,56^ and herpes simplex virus 1 ^57^. Interestingly, HSV-1 this herpes virus has a high incidence in isolated Amerindians from the Peruvian and Brazilian Amazon ^58–60^, with the elevated diversity of the virus and an endemic subtype that suggest an ancient endemic infection ^61^. This result is consistent with the hypothesis of CD45 evolution driven by the host-virus arms race model ^62^.

In conclusion, in Western South America, there is an environmental and cultural differentiation between the fertile north of the Andes, where altitudes are lower; and the arid south of the Andes, where these mountains are higher, defining sharp environmental differences between the Andes and Amazonia. This has influenced the genetic structure of Western South Amerindian populations. Indeed, the between-population homogenization of the central-southern Andes and its differentiation with respect to Amazonian populations of similar latitudes do not extend northward. Longitudinal gene flow between the northern Coast of Peru, Andes, and Amazonia accompanied cultural and socioeconomic interactions revealed by archeology, but in the central-southern Andes, these mountains have acted as a genetic barrier to gene flow. We provide new insights on the dynamics of the genetic homogenization between the populations of the Arid Andes, which are not only due to migrations during the Inca Empire or the subsequent colonial period, but started at least during the earlier expansion of the pre-Inca Wari Empire (600-1000 YBP). This evolutionary journey of Western South Amerindians was accompanied by episodes of adaptive natural selection to the high and arid Andes vs. the low Amazon tropical forest: *HAND2-AS1* (related with cardiovascular function) and *DUOX2* (related to thyroid function and innate immunity) in the Andes. In the Amazon forest environment, the gene encoding for the protein CD45, essential for antigen recognition by T/B lymphocytes and viral-host interactions, shows a signature of positive natural selection, consistent with the host-virus arms race hypothesis. This and other studies, continue to show how Andean highlanders and Amazonian dwellers provide examples of how the interplay between geography and culture influence the genetic structure and adaptation of human populations.

## Acknowledgments

We thank the Peruvian populations for their participation. We thank the members of the Laboratório de Diversidad Genética Humana (LDGH), Mateus Gouveia, Kelly Nunes, Garrett Hellenthal, Mark Lipson, Marcia Beltrame, Fabrício Santos, Claudio Struchiner, Ricardo Santos, Luis Guillermo Lumbreras, Sandra Romero-Hidalgo, Víctor Acuña-Alonzo, and Miguel Ortega for discussions or technical assistance, the staff of the Laboratorio de Biotecnologia y Biologia Molecular of Instituto Nacional de Salud (Peru) for collaborating with the Peruvian Genome Project and conducting the genotyping. Funding support: This work was supported by the Peruvian National Institute of Health (INS), the Brazilian Conselho Nacional de Desenvolvimento Científico e Tecnológico (CNPq) and the Coordenação de Aperfeiçoamento de Pessoal de Nível Superior (CAPES, programs PROEX and PRINT). V.B. is a CAPES/PEC-PG fellow (88882.195664/2018-01). Data sets were processed in sagarana HPC cluster from the Centro de Laboratórios Multiusuários from the ICB-UFMG.

