## Supplementary Text and Figures for "The genetic structure and adaptation of Andean highlanders and Amazonian dwellers is influenced by the interplay between geography and culture"

### Supplementary material

#### Summary methods

We genotyped 289 present-day Native Americans from Peru using the Human Omni array of Illumina for 2.5 million of SNPs as part of the Peruvian Genome Diversity Project. Quality control was performed using PLINK [1] and scripts *in house* [2]. We merged our individuals with public datasets [3–6] and Kaqchikel individuals from Michael Dean Lab from NCI. To generate masked data, we phased our datasets with shapeit2 [7] and inferring the non-Native DNA segments with RFmix [8]. To infer population structure we used two approaches: 1) principal component analysis in Eigenstrat [9] and genetic clustering on ADMIXTURE software [10] using an LD pruned dataset and 2) fineSTRUCTURE [11], MIXTURE MODEL [12,13] and SOURCEFIND [14] for haplotype-based analyses, after phase inference. Historical relationships were inferred using *D*-statistics [15], outgroup $f_3$ [16] and Admixture Graphs [16]. IBD and effective population size inference was done using refinedIBD [17] and IBDNe [18], respectively. For the genetic differentiation analyses, the pairwise genetic distances (F-statistics) between Native South American groups ($F_{ST}$) and between populations within groups ($F_{SC}$) were calculated for multi-locus and individual loci through 4P software [19] and hierfstat R package [20] respectively. The linkage disequilibrium was inferred by the software Haploview [21]. Natural selection scans were performed using Population Branch Statistic [22,23] and xpEHH from the package Selscan [24,25].

#### Summary methods on each data set

| The genetic structure and adaptation of Andean highlanders and Amazonian dwellers is influenced by the interplay between geography and culture | | | | |
|---|---|---|---|---|
| **Main Topics** | **Methods** | **Datasets** | | |
| | | *Natives 1.9M* | *Natives 500k* | *Natives 230k* |
| | | 1,945,113 SNPs | 515,412 SNPs | 235,405 SNPs |
| | | 384 individuals | 558 individuals | 1,047 individuals |
| Population Structure | ADMIXTURE and PCA | X | X | X |
| | fineSTRUCTURE | X | X | X |
| | MIXTURE MODEL / SOURCEFIND | X | X | X |
| Cultural interaction in a broad and regional scale were accompanied by gene flow events in Western South America | D statistics | | X | |
| | Admixture graphs | | X | |
| Dating gene flow events | RefinedIBD | X | X | |
| | IBDne | X | | |
| | Outgroup f3 | | X | |
| Genetic Differentiation and Natural Selection in Andes and Amazon | F Statistics | X | | |
| | EHH Based | X | | |
| | PBS | X | | |

**Sampling and quality control** 4

**Merging datasets** 5

**Masking Non-Native American ancestry** 6

**Section 1: Population structure of South American Natives** 7
Introduction 7
Methods 7
Genotype based methods 7
Natives 1.9M dataset 8
Natives 500K dataset 8
Natives 230K dataset 9
Haplotype based methods 10
Natives 1.9M dataset 11
Natives 500K dataset 11
Natives 230K dataset 11
Ancestry profiles 12
Natives 1.9M dataset 12
Natives 500K Dataset 13
Conclusions 14

**Section 2: Cultural interactions were accompanied by gene flow events across the Andes** 15
Introduction 15
Methods 15
D statistics 15
Natives_500K Dataset (Masked) 16
Natives_230K (Masked) 17
Admixture graphs 17
Conclusions 17

**Section 3: Dating gene flow events** 18
Introduction 18
Methods 19
Identical-by-descent segment analysis 19
RefinedIBD 19
Natives 1.9M dataset 20
Natives 500K dataset 20
IBDne 21
Natives_1.9M dataset 21
Outgroup f3 statistics 21
Conclusions 22

**Section 4: Genetic Differentiation and Natural Selection in the Andes and Amazon** 23
Introduction 23
Methods 23
Natural Selection Candidate SNPs 23
Population Branch Statistic 23
XP-EHH: cross population extended haplotype homozygosity 25
Conclusions 26

### Sampling and quality control

The present work is part of a main project called "**ANÁLISIS DE LA VARIABILIDAD GENÉTICA EN DIFERENTES POBLACIONES MESTIZAS Y NATIVAS DEL PERÚ COMO BASE PARA EL DESARROLLO DE LA MEDICINA GENÓMICA EN EL PERÚ"** or the Peruvian Genome diversity Project (PGDP), of the Peruvian Institute of Health (Instituto Nacional de Salud - INS). This is a genomic initiative to explore the genetic composition of native and admixed Peruvians. The protocol for this study was approved by The Research and Ethical Committee (OI003-11 and OI-087-13) of the INS. The PGDP was funded by the Ministry of Health of Peru and involves the collaboration of Tarazona-Santos's Laboratory of Human Genetic Diversity (LDGH) of Universidade Federal de Minas Gerais and INS. Fifteen Native American populations were sampled as part of the PGDP including populations from South Pacific Peruvian Coast (SPPC), Andean and Amazon regions (Table S1 and Figure 1).

Populations from SPPC region were sampled in northwestern Peru and involve two native communities: Moche and Tallanes. In the Andes, four communities were sampled, two Quechua-speaking communities, Qeros and Chopccas, from South Central Andes, and two Aimara-speaking populations, Jaqarus and Uros. The Amazon populations were sampled in two different ecological areas, Amazon Yunga and lower Amazon. The Amazon Yunga correspond to a cloudy forest which is a transition between the Andean mountains and the Lower Amazon. Six populations were sampled from Amazon Yunga: Chachapoyas, Lamas, Awajun, Candoshi, Ashaninkas and Matsiguenkas. Chachapoyas comprises several groups that played an important role for the Andean people as an open door to Amazon resources [26]. The Lamas population, that lives in the Upper Huallaga river, started as a "reduction" (a forced concentration of Native American groups) during the XVIII century [27]. Currently, Both Chachapoyas and Lamas groups speak Quechua. The Awajun population belongs to the Jivaroan linguistic family and its presence in the Amazon Yunga is dated back to around 1,200 years ago [28,29]. This Jivaroan population was involved in several cultural trades with South Pacific Coast populations from Peru and Ecuador but with conflicted interaction with the Inca empire (Andes). The Candoshi group is settled along the tributaries of the Pastaza River and its language has a controversial origin by being considered by some scholars as part of the Jivaro group or even independent [30]. For the Arawakan linguistic group, three individuals were collected in the Sepahua district from Ucayali region that belong to the Matsiguenkas tribe. For the Lower Amazon, three populations of the Panoan linguistic family were collected: Shipibo-Conibo, Matses and Nahuas. Finally, all native participants were required to be over 18 years old, for whom all four grandfathers were born in the selected ancestral native population.

A total of 289 individuals were genotyped using Illumina Human Omni array 2.5M at the INS. The total number of genotyped SNPs was 2,391,739. Quality control was performed using the PLINK 1.7 software [1] and in-house scripts [2]. We removed SNPs and individuals with high level of missing data (>10%), loci with 100% of heterozygous, non chromosomal information and A/T-C/G genotypes. We merged the INS data with other dataset of 127 individuals from four Native American populations from LDGH (Table S1 and Figure S1). The LDGH dataset includes four populations: two Andean (one Quechua and one Aimara-speaking) and two Amazonian (Ashaninkas and Matsiguenkas from Shimaa community). The Quechua-speaking population was sampled in a large area comprising two continuous regions, Ayacucho and Apurimac, for this reason these individuals were grouped as Quechuas_AA. In the case of the Aimara-speaking population, individuals were collected near the Titicaca lake shore in Puno region. The two Amazonian populations inhabit the Amazon Yunga area and belong to the Arawakan linguistic family. The merged data, INS and LDGH individuals, contain a total of 2,077,858 SNPs for 418 individuals organized in a total of 19 populations. Both groups, INS and LDGH datasets, include independent samples of Ashaninka population from the same region, for this reason we merge these individuals in a unique Ashaninka sample and a total of 18 populations.

Before filtering by relatedness, we removed SNPs that were in high linkage disequilibrium (LD) for each population, as it affects the inferences of relatedness, with PLINK 1.7 using the flag --indep-pairwise with the following parameters: 200 25 0.4. The first parameter indicates a window of 200 SNPs, the second indicates that the windows were advanced by 25 SNPs each time and the third indicates the LD threshold ($r^2$).

Family structure is recognized to affect the analysis of population structure as a familiar cluster can be confounded with a discrete population [31]. To overcome this issue, we estimated the kinship coefficients (Φij) for each pair of individuals for each population using autosomal SNPs. The kinship coefficient is the probability that two alleles at a locus randomly picked from individuals i and j, are identical-by-descent (IBD). For each population, we estimated the kinship coefficient using the option –genome in PLINK 1.7. We considered a thresholds of $\Phi ij \geq 0.25$ to define relatedness or not. A pair of individuals with Φij above 0.25 is defined as first-degree relatives (Parent-offspring pair and full sibling). We used a network approach to identify which individuals should be removed preserving a maximum number of unrelated individuals [32]. After applying the kinship filter, we kept 358 individuals (Table S1) for an unrelated dataset (UDataset).

### Merging datasets

We merged the UDataset with the following datasets:

- 1000 Genomes project [33].
- Human Genome Diversity Project (HGDP) [34].
- Native Americans previously genotyped by Reich *et al.* (unmasked data) [4].
- Native individuals from Guatemala (Kaqchikel population) from Michael Dean-Lab (National Cancer Institute).
- Native American individuals from two public datasets (Simons Genome Diversity Project [3] and Raghavan *et al.,* 2015 [5]).
- Ancient DNA individuals from South America [6]

From 1000 Genomes Project, we selected individuals of European (IBS, CEU), African (YRI and LWK) and East Asian (CHS, CDX, CHB) ancestries. The unmasked dataset from Reich *et al.* [4] included individuals from HGDP: Yakut, Karitiana, Surui, Pima, Maya, Piapoco, Papuan and Melanesian. From the Simons Genome Diversity Project and individuals generated by Raghavan *et al.* [5], we included all Native Americans. The available dataset of Raghavan *et al.* [5] included the ancient genome of Anzick-1 individual from Clovis complex (hereafter Clovis). Before merging individuals from Reich et al [4], we applied a relatedness filter. We removed 58 individuals with kinship coefficient above 0.1 using the same procedure employed for our samples. We generated three datasets (Table S1-S3, Figure S1) considering the density of SNPs and sample size:

a) **Natives 1.9M Dataset (**1,927,769 SNPs/623 individuals**):** Dataset with maximum number of genotyped SNPs. This dataset includes just Peruvian Native individuals from INS and LDGH and 50 Iberian (IBS), 50 Yoruba (YRI) and 50 East Asian individuals (CDX) from 1000 Genomes Project (Table S1).
b) **Natives 500K Dataset (**567,718 SNPs/773 individuals**)**: This dataset includes individuals from **Natives 1.9M Dataset,** Native American, Siberian, and Oceanian (Bougainville and Papuan), individuals from the Simons Project [3], Raghavan *et al.,* 2015 and 79 individuals from Guatemala of Michael Dean NCI lab (Table S2) genotyped for 600K SNPs. Guatemalan individuals includes individuals from Kaqchikel native population and non-native individuals with more than 99% of Native American ancestry.
c) **Natives 230K Dataset** (235,352 SNPs/1,286 individuals): Dataset with maximum number of individuals. This data include individuals from Natives 1.9M dataset and all Native Americans from the unmasked data of Reich *et al.* (2012) (~300K SNPs), which includes HGDP individuals (Table S3).

The East and South Asian, Siberian and Oceanian populations and ancient individuals from the Natives 500K and Natives 230K datasets were used only for population history analysis of the masked data and genotype based methods and not for the population structure analyses in order to avoid any confounding signal.

### Masking Non-Native American ancestry

The study sample could contain individuals some level of Non-Native American ancestries (Post-Columbian admixture). In order to infer events related to Pre-Columbian times or involving just Native American ancestry, we identified regions of European and African ancestries in Native American genomes using RFMix software [8] and then mask it. Using masked datasets and methods based on allele frequency correlation, we inferred genetic affinity among South American Natives.

Briefly, RFMix identify regions of a specific ancestry in the genome of admixed individuals using reference panels of individuals of European, African and Native American ancestries. For this purpose, we used the phased **Natives 500K** and **Natives 230K datasets**. We masked these datasets because they contain a more diverse set of populations. For the reference panels, we used 100 African (YRI and LWK) and 100 European (CEU and IBS) individuals from 1000 Genomes project as proxies. For the Native American reference panel, we selected individuals with less than 0.002% of Non-Native ancestry (European + African ancestries) using the ADMIXTURE results (see Section 1) for 3 ancestry clusters (K=3). All other Native American individuals that have some level of European or African ancestry were used as targets. We ran RFMix with the option PopPhased to enable the phase correction option. We also used two rounds of the expectation-maximization (EM) algorithm. All other settings were used as default. After run RFMix, we used the forward-backward probability output to set all local ancestry inferences that have less than 0.95 posterior probability of being Native American as missing data. Finally, the genomic regions in each sample that did not contain homozygous high quality Native American ancestry inferences were set as missing data.

### Section 1: Population structure of South American Natives

#### Introduction

The peopling and historical relationships of South American Native populations remains obscure. Previous studies based on uniparental and autosomal markers [35–37] have shown that populations living on the Western slope of the Andes, basically Andean populations, have a different pattern of genetic structure compared to populations from the Eastern slope. Based on genome-wide information, some studies have demonstrated that this dichotomic pattern of differentiation (Western-Eastern) is persistent, but with some cases of gene flow between Andes and Amazon [4,38,39]. However, Barbieri *et* al. [40] showed that some northern Peruvian populations share some ancestry cluster patterns. In this sense, our **first goal** is to explore the population structure and divergence in Western South American Natives.

To address this issue, we performed allele (ADMIXTURE and PCA) and haplotype based (CHROMOPAINTER and fineSTRUCTURE) methods. We did not perform haplotype based methods on the masked data because the masking could mislead the pattern of haplotype inference.

#### Methods

##### Genotype based methods

To understand the present-day population structure, we applied genetic clustering analysis and Principal Component Analysis (PCA). For the genetic clustering analysis, we ran ADMIXTURE [10], a population genetics model-based approach to estimate ancestry profiles. The ADMIXTURE algorithm assumes that the genetic composition of each individual is made up of up to K parental populations or ancestry clusters, where K is defined by the user. ADMIXTURE estimates the fraction that each K population contributes to an individual, as well as the allele frequencies of each of the K populations, by fitting the Hardy Weinberg equilibrium in each of the K populations/clusters. We ran ADMIXTURE in unsupervised mode for different values of K and used a cross validation (CV) test to determine the K value with the best model fitting. The ADMIXTURE results are represented as a bar plot where each individual is represented by a bar in which each color corresponds to the ancestry proportion of a specific cluster.

The PCA is a non-model based method that reduce a complex data (i.e. genotypes and individuals) to few dimensions. In the particular case of genotyping data, we can interpret these dimensions as axes of continuous genetic variation [9].

ADMIXTURE analysis and PCA assume independence among SNPs, for this reason we pruned all datasets for linkage disequilibrium (LD). We removed highly linked SNPs using PLINK 1.7 with the option indep-pairwise 200 25 0.4 for each dataset. We generate three datasets pruned by LD:

- **Natives 1.9M dataset_LDpruned**
- **Natives 500K dataset_LDpruned**
- **Natives 230K dataset_LDpruned**

We ran 50 replicates of ADMIXTURE in unsupervised mode with different random seeds for each K value and calculated the cross validation error for each run. We ran ADMIXTURE considering from K=2 ancestral clusters until the value of cross validation error starts to increases for each dataset. We plot all ADMIXTURE runs with the higher log likelihood for each K value. We ran the PCA using EIGENSOFT 4.21 [9] for the three LD pruned datasets.

#### Natives 1.9M dataset

ADMIXTURE results are displayed on Figure S2. The lower CV error was obtained for the run with five ancestry clusters (K=5). ADMIXTURE run K=3 stand out for clusters related to continental ancestry: Native American (green), European (IBS, red) and African (YRI, blue) clusters. This result showed some Native American individuals (Quechuas_AA, Chachapoyas and Moche populations) with European ancestry (~10%). Specifically, for the result with the lowest cross validation error (K=5), we observed the Andean populations as a homogeneous group (brown cluster). On the other hand, we observed an ancestry cluster (light green) predominant in Northern Peruvian populations that is shared between SPPC and Chachapoyas population (Amazon Yunga).

For the PCA (Figure S3), we excluded Africans (YRI) due to its high level of differentiation that masks the relationships in Native Americans. The first principal component (PC1, Variance explained=2.36%) showed an axis of differentiation between the European and Native American groups. We observed that some Andean, Moches, and Chachapoyas individuals have some degree of European ancestry. The PC2 (Variance explained=1.2%) separated Western (Andean and SPPC populations) and Eastern (Amazon) South American natives. Chachapoyas showed affinity with SPPC populations. Jivaroan populations (Awajun and Candoshi), were intermediate in the axis Western-Eastern. Furthermore, the PC2 showed a cline for the genetic diversity of the Amazon populations, from North (Matses) to South (Matsiguenkas). As in ADMIXTURE, both Matsiguenkas groups, Shimaa (Matsiguenkas 1) and Sepahua (Matsiguenkas 2), showed high genetic affinity. Other PCs only showed the iberian genetic diversity.

In summary, ADMIXTURE analysis and PCA showed high differentiation between populations within the Amazonia and high genetic affinity among Andean groups. Chachapoyas showed high affinity with SPPC populations. Moreover, North Amazon populations (Awajun and Lamas) showed intermediate genetic ancestry between the western and eastern Andean slope populations.

#### Natives 500K dataset

ADMIXTURE results are presented on Figure S4. For bar plot representation, we grouped Surui and Karitiana as Tupian. Mesoamerican individuals were divided in Guatemalan and Mexican (Mixe, Mixtec, Pima, Zapotec and Mayan), and we grouped the Clovis individual, two Greenland, two Aleutian and two Athabascan individuals as North America. Our ADMIXTURE runs showed the lowest CV error for eight clusters (K=8). Our description was focused on patterns not observed on the 1.9M dataset for the lowest cross validation.

ADMIXTURE run K=8 showed 6 clusters associated with Native American groups: an Andean (brown), a Mesoamerican (purple), a SPPC (pink) and three Amazon related clusters (shades of green). SPPC populations showed a predominant pink ancestry that is also predominant in Chachapoyas population. The Andean populations have a predominant brown cluster. Moreover, central Andean populations (Jaqarus, Quechuas_AA and Chopccas) showed ~10% of SPPC related ancestry. Matsiguenkas individuals were observed as a highly differentiated population since it has a specific ancestry cluster (darkgreen) which is not shared with other populations of the same linguistic group (Ashaninkas). Panoan populations (Shipibo, Matses and Nahua) showed a predominant ancestry associated with Ashaninkas population. Jivaroan groups showed a specific ancestry which was predominant in Awajun population.

For the Principal Component Analysis (Figure S5), the PC1 separated Native Americans from Europeans. Some Chachapoyas, Quechuas_AA, 1 Moche, 1 Mixtec and 1 Shipibo individuals showed affinity to Europeans due to admixture. The PC2 separated Amazon from non Amazon populations, Jivaroan and Tupian individuals were observed as intermediate between these groups. The PC3 separates a group that includes Andean and Matsiguenkas individuals from other natives. PC4 showed the separation between a group including Mesoamericans and Tupian individuals from other natives. Higher PC values showed population specific differentiation and genetic variation in the IBS population.

In summary, both, ADMIXTURE and PCA support the similarity between SPPC and Chachapoyas individuals. Moreover, our results showed the close genetic similarity between Northern Peruvian (SPPC, Awajun, Candoshi, Lamas) and Tupian with Mesoamericans. Awajun and Candoshi showed to be intermediate between Andean-Amazon axis of genetic diversity.

##### Natives 230K dataset

The following description was focused on clusters related to South American natives. The lower CV error was obtained for 18 ancestral clusters (K = 18, Figure S6B). The ADMIXTURE plots (Figure S6A) from K=3 to K=5 inferred continental ancestry clusters. The ADMIXTURE plot K=5, five ancestry clusters related to each continental region were identified: Africa, Europe, Asia, Oceania (light purple, pops 90-91) and America. ADMIXTURE K=6 showed a cluster (dark green) associated to Arawakan groups (Ashaninkas and Matsiguenkas). ADMIXTURE K=8 identified a cluster (light pink) for Costa Rican natives (Bribri, Cabecar, Chorotega, Guaymi, Huetar, Maleku, Teribe). ADMIXTURE K=9 inferred an ancestral cluster (black) related to Mesoamericans, predominantly in Pima. ADMIXTURE K=10 showed an ancestry cluster (light green) related to Awajun population which is represented most of non Andean natives. ADMIXTURE K=12 identified an ancestry cluster (gray) associated to Tupian populations (Surui and Karitiana) also observed in Mesoamericans and in non Andean natives. Furthermore, a brown cluster is predominant in the Andean populations. ADMIXTURE K=13 showed a light blue cluster associated with Pima natives. ADMIXTURE K=15 showed an ancestry cluster, darkgreen and forest green, for each Peruvian Arawakan individuals (Ashaninkas and Matsiguenkas). ADMIXTURE K=17 detected a cluster (light blue) associated with SPPC population, and which is also present in Chachapoyas and Lamas. The ADMIXTURE K=18 with lowest cross validation showed differentiation in Asian natives. ADMIXTURE K=21 showed an ancestry cluster related to Lamas sample.

In the Principal Component Analysis (Figure S7), the PC1 showed the differentiation between the Old world from the Native American ancestry. Old world ancestry included Europeans (IBS), Oceanians, Asians (Russian and Mongolians) and admixed Native Americans. Greenland natives were shown as intermediate between the Old and the New world groups. In the Native American axis, the Matsiguenkas individuals were the most differentiated. Considering PC1 and PC2, we discriminated three blocks of Native ancestry, one Eskimo-Aleut (Greenland), the second Athabaskan and Algic (North American group), and the third including all other Native Americans. This pattern is consistent with Reich *et al.,* (2012).

In summary, the Andean region keeps its high homogeneity. The ancestry profile of SPPC and Chachapoyas natives was similar. Moreover, ADMIXTURE and PCA showed some genetic affinity between Northern Peru, Northern South America and Mesoamerican populations. In addition, PCA and ADMIXTURE results for masked and unmasked datasets have showed similar patterns: Andean homogenization, the close relationship between SPPC populations and Chachapoyas, Awajun as intermediate of Western and Eastern ancestries, and genetic affinity of Mesoamericans with SPPC and some Amazon populations.

#### Haplotype based methods

To get deep in the population structure of Peruvian natives we used haplotype-based methods (CHROMOPAINTER and fineSTRUCTURE algorithms), which explore the patterns of haplotype similarity among individuals [11]. First, we phased our datasets using shapeit2 software [7]. For the phasing process, we used the complete dataset (without LD pruning). To increase the accuracy of the phasing process, we used 200 conditioning states and 30 main iterations of MCMC.

The haplotype-based methods are based on the identification of the LD patterns along the genome of individuals in order to infer the number and length of DNA chunks (CHROMOPAINTER) shared among them. Then, this information is exploited in the identification of cluster of individuals based on the pattern of genetic similarity at a fine scale (fineSTRUCTURE). The process of identification of LD patterns is called chromosome painting and is performed by CHROMOPAINTER [11]. In this process each haplotype (recipient) is reconstructed with donated (or shared) chunks from other individual haplotypes (donors) [11]. For this inferences CHROMOPAINTER requires phased data and two scalar parameters (inferred in a previous CHROMOPAINTER run): 1) the recombination scaling and 2) mutation parameters. The result of the chromosome painting is summarized in two interindividual matrices called coancestry matrices. These matrices explain donor-recipient relationship as the total number (chunkcounts) and the length (chunklengths) of DNA chunks shared among individuals.

After the chromosome painting, we used the chunkcounts co-ancestry matrix to infer the population structure using the model-based approach fineSTRUCTURE [11]. If two individuals belong to the same population (here inferred as a cluster), we expect a similar distribution of sharing chunks summarized in the co-ancestry matrix. Using a reversible-jump MCMC, fineSTRUCTURE assigns individuals into clusters that may resemble their populations. Like other MCMC algorithms, fineSTRUCTURE is dependent on the number of MCMC iterations so it uses a previous burn-in stage and then for several iterations (i.e 2 millions).

Since our main question is about the history of Native Americans, we excluded individuals with high levels of Non-Native ancestry (>5%) in the Natives 230K dataset except for Chachapoyas individuals. We did this because Chachapoyas population have almost all individuals with more than 5% of Non-Native ancestry, so we maintained all individuals (see

ADMIXTURE results). We removed individuals just in this dataset because it has a better representation of Native Americans. We determined which individuals had to be removed based on ADMIXTURE results (K=3). For Natives 1.9M and Natives 500K datasets we maintained the complete dataset.

Using the Expectation-Maximization algorithm for a subset of individuals and chromosomes for each data set, we obtained the following values for the parameters. The recombination scaling and mutation were respectively: 220.324 and 0.00018 for the Natives 1.9M dataset, 144.362 and 0.0002 for the Natives 500K dataset, and, finally, 150 and 0.00045 for the Natives 230K dataset. After obtaining the co-ancestry matrix, we ran fineSTRUCTURE for all datasets considering 1,000,000 of burn-in steps, 2,000,000 of MCMC iterations and 100,000 of sampling. After the MCMC calculations, we construct the tree using 100,000 additional steps of hill-climbing steps. We represented the fineSTRUCTURE results as a tree and the chunklengths coancestry matrix as a heatmap.

##### Natives 1.9M dataset

The fineSTRUCTURE tree clusterize the native individuals in three main groups: Admixed natives (Figure S8A), Amazon (Figure S8B), SPPC and Andean individuals (Figure S8C). Almost all Native Americans were grouped in clusters containing individuals of the same population label. Most of the SPPC individuals showed genetic affinity with Andean populations. Arawakan groups (Ashaninkas and Matsiguenkas) showed higher genetic affinity, the same observation is for Panoan groups (Matses, Nahua, Shipibo) but no for Jivaroan groups (Figure S8B). Chachapoyas population showed a close relationship with the admixed Native Americans.

##### Natives 500K dataset

The arrangement of the clusters was similar to the resulting tree of Natives 1.9M dataset with some novelties (Figure S9): 1) Moche and Chachapoyas individuals showed close relationship with Andean individuals (Figure S9A), 2) West Mesoamerican natives (Mixe, Mixtec, Zapotec and Pima) clustered together with some Amazon Tupian (Surui and Karitiana) individuals (Figure S9B), 3) Peruvian Arawakan natives (Matsiguenkas and Ashaninkas) were shown as the most differentiated clusters (Figure S9D).

##### Natives 230K dataset

The fineSTRUCTURE tree (Figure S10A and B) showed a more external cluster that contains IBS, Chipewyan and Greenland natives. The second more external clusters include Arawakan speakers as the most differentiated populations (Figure S10B). Other Native Americans were organized in two macro clusters, Andean (Figure S10B) and non-Andean clusters (Figure S10). The Andean cluster (Figure S10B) showed no major differences from the clustering of the other datasets, showing Uros as the most differentiated Andean population. Costa Rican populations (Figure S10B) showed close affinity to Northern South American populations (Embera, Wayuu, Waunana and Kogi). Moreover, Mayan populations (Mayan and Kaqchikel individuals) showed close affinity to Northern Peruvian (Lamas and SPPC) and Inga individuals (Figure S10A). Panoan individuals (Matses and Nahua) clusterize with other Eastern populations (Figure S10A).

Summarizing the fine-scale population structure analyses, we inferred the close relationship of SPPC natives with Mesoamericans (Maya and Kaqchikel) and a relationship between SPPC individuals and Chachapoyas.

#### Ancestry profiles

Native American populations were affected by several episodes of genetic drift which can mislead the ADMIXTURE and CHROMOPAINTER/fineSTRUCTURE interpretations. Drifted populations tend to generate ancestry clusters in the ADMIXTURE analysis. The Chromosome painting inference using the "-a" switch (employed for fineSTRUCTURE analyses), could be problematic for higher drifted populations because its individuals tend to copy more times from individuals of the same group than from other populations. Because of this, we cannot have a clear pattern of similarity among native populations.

To overcome this problem, we performed a Chromosome painting inference for each Native American population setting a population as recipient from all other populations (CHROMOPAINTER "-f" switch). This inference result in a chunklengths matrix that summarizes the contribution (shared DNA) from the donor populations to the recipient. Then, with this matrix, we applied two approaches to infer the ancestry proportions: a regression model (non-negative least squares) implemented on GLOBETROTTER software (MIXTURE MODEL) [12,13,41] and a Bayesian model implemented in SOURCEFIND [14]. To generate the chunklengths output for each Native American groups, we used the same two scaling parameters inferred in the last subsection. Furthermore, since fineSTRUCTURE and ADMIXTURE showed no differences among Matsiguenkas individuals, Matsiguenkas 1 (Shimaa) and Matsiguenkas 2 (Sepahua), for GLOBETROTTER inferences we considered these two groups as one single Matsiguenkas group.

##### Natives 1.9M dataset

The MIXTURE MODEL (Figure S11) revealed a particular pattern in Andean populations. Each Andean population share a high proportion of DNA with other Andean populations showing a homogeneous pattern. Specifically, the ancestry profile of Uros population showed that 95% of its DNA is shared with Puno population indicating lower genetic variability of Uros. Moreover, Aimara-speaking Jaqarus showed high genetic affinity with Quechua-speaking groups. The SPPC populations have more affinities with northern amazon populations. These affinities are related to the similar ancestry proportions and the ancestry related to Chachapoyas. Also, the three North Amazon populations (Candoshi, Awajun, Lamas and Chachapoyas) showed a significant proportion of ancestry related to SPPC populations (>10%). Furthermore, we observed that SPPC and North Amazon populations have a significant proportion of shared DNA with Andean population (>20%). SOURCEFIND tends to eliminate contributions that could be due to background noise. For this particular dataset, SOURCEFIND identify the SPPC ancestry in north Amazon populations and that most of the Andean ancestry is explained by Quechua related ancestry (Figure S11). For this dataset, it is important to consider the poor representation of other native populations, Mesoamericans and South Americans. This poor representation gave us two interpretations to the Andean shared ancestry in SPPC and North Amazon populations: 1) real Andean ancestry or 2) In the absence of a closely related ancestry, the andean ancestry fill this vacuum. The Amazon populations showed a common pattern of genetic composition among them. Only Quechuas_AA, Jaqarus, Moche and Chachapoyas showed European ancestry.

#### Natives 500K Dataset

The MIXTURE MODEL (Figure S12) showed the Andean populations with the same pattern as was observed with the Natives 1.9M dataset, they share a high proportion of DNA with other Andean populations showing a homogeneous pattern. Furthermore, it is possible to observe Mesoamerican related ancestry in almost all Peruvian natives. The ancestry composition of SPPC populations, showed more similarity to Chachapoyas population. Moreover, Jivaroan and Lamas populations showed very similar patterns of ancestries. We observed that Moche and Chachapoyas populations have a significant proportion of shared DNA with Andean population (~30%). All SPPC and Amazon populations (except Ashaninkas and Matsiguenkas) showed more ancestry related to Mesoamerican (>18%) than to the Andean populations. Matsiguenkas showed more ancestry related to Ashaninkas, indicating a close genetic affinity among Arawakan populations. SOURCEFIND showed that the Mesoamerican related ancestry is particularly high in SPPC and Karitiana populations. The Andean contribution in the Amazon was reduced and restricted to Chachapoyas population. Moreover, Jivaroan (Awajun and Candoshi) populations showed some contribution from SPPC group.

#### Natives 230K Dataset

Using fineSTRUCTURE results (Figure S10A and B), we organize Native American populations in the following clusters:

- Chopccas (17)
- Quechuas_Per (13) [Quechuas_AA and Quechua_R2]
- Aimaras_PB (29)
- Qeros (12)
- Uros (13)
- Quechuas_Bol (10)
- Ashaninkas (35)
- Matsiguenkas (26)
- Matses (11)
- Nahua (2)
- Awajun (23)
- Lamas (21)
- Chachapoyas (9)
- Moche (25)
- Tallanes (34)
- Pima (21)
- Tepehuano (21)
- Maya (29)
- Kaqchikel (6)
- Mixe (17)
- Zapotec1 (6)
- Zapotec2 (21)
- Mixtec (5)
- Karitiana (8)
- Surui (10)
- Guahibo (6)
- Palikur (3)
- Piapoco (6)
- Ticuna (4)
- Embera (4)
- Kogi (3)
- Waunana(3)
- Wayuu (2)
- Toba (4)
- East_Amazon_Brazil [ Arara (1) - Parakana (1)]
- Inga (2)
- 1 Chaco cluster [Guarani (3) - Chane (2)]
- Wichi (4)
- Maleku (2)
- Cabecar (16)
- Guaymi (5)
- Teribe (3)
- Chipewyan (3)
- East Greenland (3)
- IBS cluster (107)
- CDX cluster (93)

The number in parenthesis indicates the number of individuals after filtered the ones with more than 5% of European ancestry. Also, we did not include Chono and Huilliche as donors due to the low probability that they were involved in gene flow events with Central Andes or Amazonia [4,42]. Furthermore, the Quechuas individuals from our dataset that clustered with Quechuas R2 from Reich *et al.* [4] were included as Quechuas Per and all Quechuas R1 were considered as Quechuas Bol (from Bolivia). Although Jamamadi sample form a cluster with Arara and Parakana, these individuals are geographically distant from Arara and Parakana, for this reason we excluded them from the GLOBETROTTER analysis. To improve the visualization of the results, we grouped the following populations:

- North Amazon 1: Inga and Ticuna.
- North Amazon 2: Guahibo, Palikur and Piapoco.
- Caribbean: Embera, Kogi, Wayuu and Waunana.
- Chaco natives: Wichi, Guarani and Chane.
- Pampas: Toba.
- Central America: Cabecar, Maleku, Guaymi and Teribe.
- Mayan : Maya1 and Maya 2.
- West Mesoamericans: Mixe, Mixtec, Zapotec1 and Zapotec2.
- North American: East Greenland and Chipewyan.

The MIXTURE MODEL (Figure S13) showed a contrasting pattern between Western and Eastern slope populations. The SPPC populations as well as Amazonian groups showed contribution from Mesoamerican groups. SOURCEFIND showed that the mesoamerican contribution is restricted to SPPC, Chachapoyas, Awajun and Inga groups. Moreover Chachapoyas and Inga groups showed some level of Andean related ancestry.

### Conclusions

- The genetic dichotomy between populations living on the Western and Eastern slope of the Andes does not extend to the Northern Peruvian populations.
- Chachapoyas, a population living in the Eastern slope of the Andes, have higher genetic affinity with SPPC populations.

### Section 2: Cultural interactions were accompanied by gene flow events across the Andes

#### Introduction

Archaeological evidence suggest an intensive interaction involving South American natives from different regions (i.e. Andes and Amazon). In a regional scale, cultural and commercial interactions between South Pacific Coast, Andean and Amazonian populations are well documented [28,29,43,44]. Archaeological data point out that the region involving the Southern Ecuador and Northern Peru was a main crossroads for the Andes-Amazon interaction [43], which could be facilitated since the Andean mountain chain has its lowest altitude in this region [45,46]. These cultural and commercial interactions involved the trade of Spondylus shells from the Coast and medicinal plants and herbs from the Amazon. Particularly, Chachapoyas, a population that inhabit this area, was part of a significant interregional exchange network during the Early Intermediate period (around 2300 YBPto 1400 YBP) and that extended to Inca period (around 1500 AD) [26]. Also, it was suggested a socioeconomic exchange between Moche (Coast) and Awajun (Amazon Yunga) natives [28]. Nowadays, Chachapoyas and Lamas populations both living in the Amazon Yunga speak Quechua, that could be adopted as a lingua franca in the last centuries. Specifically, Lamas population is an intriguing case since was attributed to have an Andean Origin (related to Chanka population). A recent study demonstrated that Lamas population has a closer genetic affinity to surrounding Amazon populations than with Chanka population, suggesting a possibly Amazonic origin instead of Andean [47]. In the latter section, we showed the genetic affinity among Northern Peruvian populations. This genetic affinity could be related to the historical interactions. With new genome-wide data from populations from Amazon Yunga and SPPC, our **second goal** is to explore **whether gene flow accompanied the cultural and socioeconomic interactions between Andean and Amazon Yunga populations** or not. To address this question we performed *D* statistics and Admixture graphs analyses on the masked data. The *D* statistics determine if two populations have an excess of alleles sharing due to gene flow. Moreover, the admixture graphs explore the best model of relationships among populations taking into account gene flow events.

#### Methods

##### *D* statistics

The *D* statistics [15,48], or BABA-ABBA test, is a method to detect secondary contact among closely related populations. It evaluates four populations: $P_1$, $P_2$, $P_3$ and an outgroup:

$$Outgroup\ (P_3\ \ (P_1,\quad P_2))$$

This treeness test considers, as a null hypothesis, that $P_1$, $P_2$, $P_3$ and an outgroup have a tree relationship. In this null hypothesis, $P_1$ and $P_2$ diverged earlier from the ancestor of $P_3$, with no gene flow between $P_3$ and $P_1$ or $P_2$ after the divergence. The alternative hypothesis is that $P_3$ was involved in gene flow with $P_1$ or $P_2$ after the divergence. This analysis is restricted to biallelic sites and considered an allele "A" as the ancestral allele of the outgroup (O) and an alternative allele “B” in $P_3$. Considering the order $P_1$-$P_2$-$P_3$-O, the *D* test is focused on the BABA or ABBA pattern. The first pattern (BABA) correspond to the total number of sites with the alternative allele (B) shared only by $P_1$ and $P_3$. The second configuration (ABBA) is the total number of sites with the alternative allele (B) shared only by $P_2$ and $P_3$. The *D* statistic is calculated by the relationship of: the difference of BABA-ABBA counts (numerator) and the total count BABA+ABBA (denominator). The numerator of this relationship indicates the signal of the statistic which is interpreted as the direction of gene flow. If the *D* statistic is not significantly different from zero, we accept the null hypothesis of treeness. If the result differ significantly from zero, we reject the null hypothesis and consider the possibility of gene flow between $P_3$ with $P_1$ or $P_2$. For *D* statistics estimation we used ADMIXTOOLS [16]. The results are interpreted as follows: negative values of *D* are interpreted as secondary contact between $P_2$ and $P_3$ and, positive values indicate a secondary contact between $P_1$ and $P_3$. A *D* value is considered statistically significant if the absolute value of the relationship between the *D* value and the standard deviation is equal or above 3 (|Z score| □3). We applied *D* statistics to the masked Datasets 500K and 230K.

Considering the huge number of combinations for *D* inferences and even obtaining highly significant values, we construct Q-Q plots to show if these results could be expected by chance or not.

<u>Cultural interactions and gene flow across the Andes</u>

Considering the divergence between Western (including Andean, SPPC) and Eastern South American clusters, we tested the following configurations:

- (Outgroup, Eastern (Western$_2$, Western$_1$))
- (Outgroup, Western (Eastern$_1$, Eastern$_2$, ))

Outgroup: For both masked datasets, we used Africans (YRI).

##### Natives_500K Dataset (Masked)

Configuration **(Outgroup, Western (Eastern$_2$, Eastern$_1$)):**
We explored the gene flow among clusters in South America. This configuration explore if one western population shares more alleles with one population from the eastern slope than another. Deviation from the diagonal of the Q-Q plot indicates that some populations in this configuration are involved in gene flow events. We found signals of gene flow involving Amazonian populations with SPPC (Figure S14).

Configuration **(Outgroup, Eastern (Western$_2$, Western$_1$)):**
When we explored the possible signal of gene flow from Eastern (Amazon yunga and Lower Amazon) to Western, we found results similar to the first configuration, a significant deviation from the diagonal involving SPPC, with Chachapoyas, Lamas and Jivaroan populations (Figure S15).

##### Natives_230K (Masked)

For this dataset, both configurations showed congruent results with the Dataset 500K (Figures S16, S17).

#### Admixture graphs

Our population structure analyses (first section) suggest a higher genetic affinity between populations across the Andes in the fertile region (Northern Peru, SPPC and Chachapoyas) in contrast to the arid region (Andean populations). *D* statistics gave support to this idea. In order to build a model of population relationships to understand the phylogenetic relationships and gene flow events we used admixture graphs. We model the relationships using the masked Dataset 500K on a subset of populations to avoid overfitting.

The subset includes a SPPC (Tallanes), two Andean (Chopccas and Uros) and three Amazon populations (Chachapoyas, Ashaninkas and Matses). We also included two Mesoamerican (Mayan and Mixe), East Asians (CDX) and Africans (YRI). We fit a model to explain the relationships of Awajun, Candoshi and Lamas that showed significant values of D statistics.

Three admixture graphs were obtained (|Z score| < 3) that showed a highly dynamic relationship between SPPC, Andean and Amazon populations (Figures S18-S20). The first admixture graph (|Z score = 2.689|) showed that the most basal Amazon ancestor resulted of and admixture event among Mesoamerican and South American ancestor. SPPC population (Tallanes) and Chachapoyas are fitted in the middle of several gene flow events into the Amazon (Awajun) (Figure S18).
The second admixture graph explain the best fit for the Candoshi population (|Z score|= 2.801). This graph shows that Candoshi population is involved in similar patterns of gene flow like Awajun, receiving a significant contribution from western slope (20%) (Figure 19).
The third graph (|Z score| = 2.98) shows that Lamas also result from the Western slope gene flow (Figure S20).

### Conclusions

- D statistics and Admixture graphs suggest that Tallanes population was involved in gene flow events into the Amazon Yunga populations specifically Northern Amazon populations.
- Chachapoyas, Lamas, Awajun and Candoshi populations are fitted as admixed populations between an SPPC population and an Amazon ancestor with at least 20% of genetic contribution from Tallanes like population.

### Section 3: Dating gene flow events

#### Introduction

Our results in latter sections showed evidence of gene flow at two main patterns: 1) gene flow across the Andes and 2) the Central Andean genetic homogenization. In the following section, we explored IBD based methods and outgroups f3 in order to infer the time frame involving gene flow signals and demographic patterns.

Specifically, we addressed three issues:

- **Gene flow across the Andes:** The Northern Peru was characterized by the limited effect of geographical barriers for the cultural movement across the Andes. In the latter section, we found evidence of gene flow between SPPC and Northern Amazon (Awajun and Lamas) populations. Moreover, we inferred high genetic affinity for the Chachapoyas and SPPC populations. But **which was the historical context in which these genetic interactions occurred**?

- **Central Andean genetic homogenization:** The Andean archaeology has classified the last five millenia based on: 1) cultural expansion (Horizons) and 2) the falling of empires and development of local cultures (Intermediate) (Fig S21 - Scliar et al. 2014). Furthermore, during colonial times, Spaniards forced some Andean populations to migrate from the birth place in order to control them and avoid rebellions. Our results (Figure 1, S3-S7) and previous studies have shown that the current genetic make-up in the Andeans is homogeneous [35,36], but since **when this phenomenon occurred and how this pattern evolved in the last centuries?** Moreover, several estimates about the population size of Andean populations characterized a dramatic change after the European arrival [49]. Recently, Lindo et al. [39] have identified a population contraction of around 27% in the Andes highlands after European contact. But, **how has the the Andean effective population size evolved before and after the European contact?**

#### Methods

##### Identical-by-descent segment analysis

We analyzed the pattern of segments identical-by-descent (IBD) to infer the relationship among populations across the time. If two DNA segments are identical and have the same ancestral origin they are considered Identical-by-descent [50,51]. From one generation to another, large segments of DNA are inherited, but in successive generations recombination events break these regions [52]. The relationship between the size of an IBD segment found between two individuals and the time in generations until coalescence have the following approximation [53,54]:

$$E = 3/2L$$

Where:
E: time in generations to the most recent common ancestor.
L: length of IBD segments (in units of Morgan).

To infer the pattern of gene flow along the time in regional and broad scale, we ran refined IBD software [17] with the **Natives 1.9M Dataset** and **Natives 500K Dataset**. To analyze the demographic evolution in Central Andes, we used IBDne software [18] with the **Natives 1.9M Dataset**, both approaches are described below.

#### RefinedIBD

To infer IBD segments, we used RefinedIBD [17] . This software performs two steps: first, it uses the GERMLINE algorithm [55] for IBD detection, and second, a refinement step, that calculates the probability of each segment to be IBD [17]. We removed all missing data in the specific dataset selected for this analysis using PLINK (--geno parameter). We used the genetic map GRCh37 from HapMap and we restricted our analyses to segments larger than 3.2cM. We organized the IBD segments in four intervals that could be related to historical periods:

1) 3.2 to 7.8cM (47 to 19 generations before present)
2) 7.8 to 9.3cM (19 to 16 generations before present)
3) all segments greater than 9.3cM (16 generations before present to present day)

The first interval is related to pre inca times, more specifically to the Middle Horizon and Late Intermediate, that correspond to the Wari-Tiwanaku Empire. The second interval involves the rise and fall of the Inca Empire. Finally, the last interval is related to colonial times until the present day (Figure 2, Figure S21).

We calculated the average amount of shared DNA between two individuals from the same (aaIBD) or different populations (abIBD) [17]. Considering a specific pair of populations (a and b), we calculated the total amount of shared DNA between one sample from “a” and another from “b”. After that, we sum all pairwise values and divided by the number of pairs between a and b:

$$\text{abIBD} = \frac{\Sigma' ij Lij}{Npairs}$$

Where:
abIBD: average of the total shared IBD length between two individuals from different populations (or the same population if it is aaIBD).
*i* and *j*: the two individuals.
*L*: total IBD length shared between each pair of individuals
*Npairs*: Na * Nb (for different populations),and Na(Na-1)/2 (For same population). Where N is the number of individuals in the respective population (a or b).

The representation of IBD relationships was presented as a heatmap constructed with the log of the abIBD values, in which the intensity of hot colour is an indication of a close relationship (high rate of sharing IBD segments) while the cold colors indicates a more distant relationship (Figure S22-S23).

##### Natives 1.9M dataset

In the first interval (3.2 to 7.8-5 cM, Figure 2B, S22A) is possible to observe homogeneous pattern among andean populations. We did not observe differences between the intra and interpopulation sharing in the Andes. In a temporal view, this interval coincides with the Middle Horizon and Late Intermediate [56] that included the expansion Tiwanaku-Wari and its falling. This society, which dominated the political landscape of the central highlands of the Andes, was probably an ancestor of Quechua-speaking populations [57], which may be related to the fact that 3 of the 6 studied populations speak this language today. Posteriorly, the difference between intra and interpopulational sharing ratio for Andean populations gradually increased until the most recent interval, but remains smaller than other groups. However, the hypothesis that the Andean homogenization already existed before the Incas was evidenced by the visualization of the high degree of sharing of IBD segments between these groups during the Tiwanaku-Wari expansion. We also observed that the genetic affinity between Chachapoyas and Western slope populations is constant in the three intervals. Moreover, the first interval showed a major genetic connection between Andes and Amazon represented by a high level of shared ibd segments.

##### Natives 500K dataset

In the first interval (Figure S23A), the Andean region already appears homogeneous, corroborating the **Natives 1.9M dataset** results. The SPPC populations shown high internal affinity degree. The Amazon group in general have some relations with other groups, but the diagonal is very intense, evidencing its high degree of intrapopulation relations. In the second interval (Figure S23B), corresponding to the period during the Inca empire, Andes stay homogeneous. In the last interval (Figure S23C), after the Europe conquist, Andes is apparently more structured. SPPC populations remain connected since the first interval, as did Matses and Lamas. Like the first dataset, the genetic affinity between Chachapoyas and Andean and SPPC is constant along the intervals. Moreover the genetic connectivity between Andes and Amazon is observed in the first interval.

#### IBDne

To understand the demographic dynamics of Andean populations, we calculated the pattern of effective population size (Ne) with software **IBDne** [18]. This algorithm is based on the Wright-Fisher model and calculates the expected distribution of the coalescent time for an IBD-segment length. Therefore, this software equate the expected and observed amount of IBD segments in a generation "g", to infer the pattern of Ne along the generations. This method have some particularities that need to be taken into account for a reliable interpretation: 1) it tends to smooth over sudden changes in Ne, 2) it assumes a closed population, that can inflate the estimates of Ne relative, 3) it assumes a homogenous population.For this reason we performed the analysis just for the Andean group. To avoid the underestimate of effective population size, we restricted the analysis to segments larger than 4cM, as is indicated by authors for array data [18]. We inferred this parameter (Ne) only between 4 and 50 generations before the present, because segments related to the last 3 generations are not informative for the dynamic of the population. As our Andean populations are more genetically homogeneous, we grouped as a unique population for the IBDne inference, which would not be possible for the other groups. Moreover, as the density of SNPs is also an important factor for these inferences, we applied this method for **Natives 1.9M dataset**.

##### Natives 1.9M dataset

In the first heatmap interval, approximately between 47 to 19 generations before present (Middle horizon), we can see an expansion period in the population effective size, that is expected in a period commanded for a unique empire, the Wari-Tiwanaku Empire. After approximately 27 generations, the Ne continuously decreased (Figure S24), which can mean a bottleneck or continuous population structuring.Considering the lower homogenization of the Andean population in this time, shown by the heatmap (Figure S22), along the Inca Empire, the second option seems more plausible.This reduction stopped in the last 10 generations.

### Conclusions

- The Andean homogenization already existed before the Late Intermediate. Probably related to the Tiwanaku-Wari expansion.
- The dynamic of Andean Ne showed that these populations have decreased in size in the last 27 generations
- Evidence of gene flow across the Andes, between Western Slope Andean and North Amazon Yunga populations was inferred before the Inca period.
- Gene flow between Mesoamerica-South America was inferred before the Inca period and could exist in more ancient times.

### Section 4: Genetic Differentiation and Natural Selection in the Andes and Amazon

#### Introduction

The evolutionary mapping of genetic variants is an efficient approach to identify functional genomic regions that have played an essential role in survival, and possibly have consequences for human health [58,59]. The evolutionary history of modern humans is marked by major migration events for environments with different climates, diets, and diseases [60]. These factors compose the selective pressures that act on variants that affect biological mechanisms that influence the adaptation process [61,62]. The process of natural selection leaves genetic signatures that can be detected, making it possible to identify regions of the human genome related to these mechanisms.

In the following section, we applied statistical methods based on population differentiation (Population Branch Statistic - PBS) and linkage disequilibrium (cross population extended haplotype homozygosity - XP-EHH) to identify genomic regions under natural selection in Andean and Amazon populations. For this purpose, we used the **Natives 1.9M dataset** considering only the following populations organized in **two groups**:

1) **Andean group:** Chopccas, Quechuas_AA, Qeros, Puno, Jaqarus, and Uros.

2) **Amazon group:** Ashaninkas, Matsiguenkas (including Matsiguenkas 1 and 2), Matses and Nahua. We did not include Awajun, Candoshi, Lamas and Chachapoyas in this analysis because our previous results (Section 1 and 2) demonstrated that these populations were involved in gene flow and this may mask differentiation signals.

#### Methods

##### Natural Selection Candidate SNPs

###### Population Branch Statistic

PBS is a statistic test to identify changes in the allele frequencies of a target population occurred since its divergence from an ancestral population. PBS is based on the comparison of differentiation (FST) values among 3 groups: 1) the target population; 2) a population close to the target, and 3) an outgroup [22].

Before the PBS analysis we applied MAF (Minimum Allele Frequency > 0.05) filter with PLINK. Since we are searching for evidence of differentiation between Andes and Amazon, we considered only SNPs with low differentiation inside these groups ($F_{SC}$<0.15 [63]). The $F_{SC}$

for each SNP for each group was estimated with varcomp function from the hierfstat R package [20]. 4P software [19] was used to calculate $F_{ST}$ for each SNP. The F-statistics estimated through varcomp function and 4P rely on the Weir and Cockerham (1984) algorithm [64]. Subsequently, the $F_{ST}$ values were transformed as following [65]:

$$F_{ST}T = -\log(1-F_{ST})$$

To the transformed $F_{ST}$ values, we applied the PBS formula [22]:

$$PBS = (F_{ST}T1+F_{ST}T2-F_{ST}T3)/2$$

Where:

$F_{ST}T1$: transformed $F_{ST}$ between the target population and the closely related population.

$F_{ST}T2$: transformed $F_{ST}$ between the target population and the distant population.

$F_{ST}T3$: transformed $F_{ST}$ between the close population and the distant population.

To avoid spurious outliers when the branches were long or short in all groups, we applied a normalized version from PBS [23]:

$$PBSn = PBS1 / (1 + PBS1 + PBS2 + PBS3)$$

Where:

PBSn: normalized PBS.

PBS1: estimated PBS when the PBS is calculated for the target population.

PBS2: estimated PBS when the PBS is focus on the close population.

PBS3: estimated PBS when the PBS is focus on the distant population.

Our final result is based on the PBSn.

We performed PBS with the following configurations: 1) Andes as a target group, Amazon as a closely related group; and 2) Amazon as a target group, and Andes as a closely related group; in both approaches the CDX (Chinese Dai in Xishuangbanna, China), a population from 1000 Genomes [66] was used as an outgroup. We analyzed the results in windows of 20 SNPs with 5 SNPs of overlap. To determine the probability that a PBS value occurs under the null hypothesis of genetic drift, we simulated 10,000 chromosomal regions of 1Mb under the neutral model for the three populations involved (Andes, Amazon and CDX) (Figure S25) using the Recosim program to simulate the recombination maps and Cosi2 [67] to simulate the genetic data under a neutral model. After this, we estimated the PBS values for the simulated data with the same methodology used for empirical data. For each observed PBS result, we calculated the p value as a proportion of simulated PBS values that are equal or greater than the observed value [22]. We considered as candidates for natural selection those SNPs in the 0.05% higher values of PBSn (PBSn > 0.150 for the Andes and PBSn > 0.191 for the Amazon) that were encompassed in the windows in the 0.05% higher PBSn mean values (PBSn mean > 0.095 for the Andes and PBSn mean > 0.116 for the Amazon). We found 142 signals comprising 16 genes in the Andes and 137 signals comprising 15 genes in the Amazon (Tables S4, S5; Figure 3).

#### XP-EHH: cross population extended haplotype homozygosity

Positive selection events increase the frequency of a genetic variant and, consequently, the frequency of the variants around it [68]. This process occurs faster than the haplotypes are broken down by recombination, leading to the emergence of an unusual high frequency long-range haplotype. Considering this, we decided to perform a homozygous extended haplotype (EHH) test concomitantly with the Population Branch Statistic (PBS) method to select the most likely candidates for natural selection.
Sabeti *et al* [24,69] developed methods to detect natural selection signatures calculating the extended haplotype homozygosity (EHH), defined as the probability of finding homozygosity of all SNPs around the haplotype of interest choosing two random chromosomes containing this haplotype in a population:

$$EHH(x_i) = \sum_{h \in C(x_i)} \frac{\binom{n_h}{2}}{\binom{n}{2}}$$

Where $C(x_i)$ is the number of all possible distinct haplotypes considering the extension from de core SNP to the i-th SNP, and $n_h$ is the number of observed haplotypes of a specific type h [25].

XP-EHH analysis were performed with the software Selscan (Szpiech and Hernandez, 2014). We considered as positive signals for natural selection the SNPs representing the 95.5 percentile of the XP-EHH results (XP-EHH > 2.97 for the Andes and XP-EHH < -3.34 for the Amazon) and as strong candidates for natural selection only the results concordant with the PBS test. With this approach we found 22 candidate SNPs comprising 3 genes in the Andes and 21 SNPs comprising 1 gene in the Amazon (Tables S6, S7; Figure S26, S27).

The candidate loci for natural selection were annotated with MASSA (Multi-agent Annotation System) [70]. Considering a list of positive signals (dbSNP rs IDs), the MASSA algorithm provides annotations based on 11 public databases.

It is interesting to note that the strongest signal for PBS analysis for the Andean populations was from the the DUOX genes (Figure S26), previously suggested as a candidate gene for natural selection by Jacovas *et al*. [71]. However, this signal does not appear in the XPEHH results. This is probably because the XPEHH method is designed to detect almost complete hard sweeps, in which a variant under selection increases to almost fixation in a population while remaining variable in other populations [24], and this is not the case for the DUOX2 variants. In fact, these variants are present at a frequency of 0.53 in the Andean populations, while they are almost fixed in other non-African world populations (Table S4).

### Conclusions

- We have confirmed a natural selection signal from a gene previously reported in Andean populations, DUOX2 [71].
- We identified Natural selection signals in genes related to high altitude adaptation (SULT1A1, RARS) [72,73], heart development (HAND2-AS1) [74] and immune response (UBQLN4 SSR2, DUOX2) [75–77] in Andeans (Tab. S4).
- We identified Natural selection signals related immune response (PTPRC) (Meer et al. 2019), food intake regulation (MCHR1) [78] and lipid transport (ABCA9, ABCA6) [79,80] in Amazon populations (Tab. S5).

**Supplementary Figures**

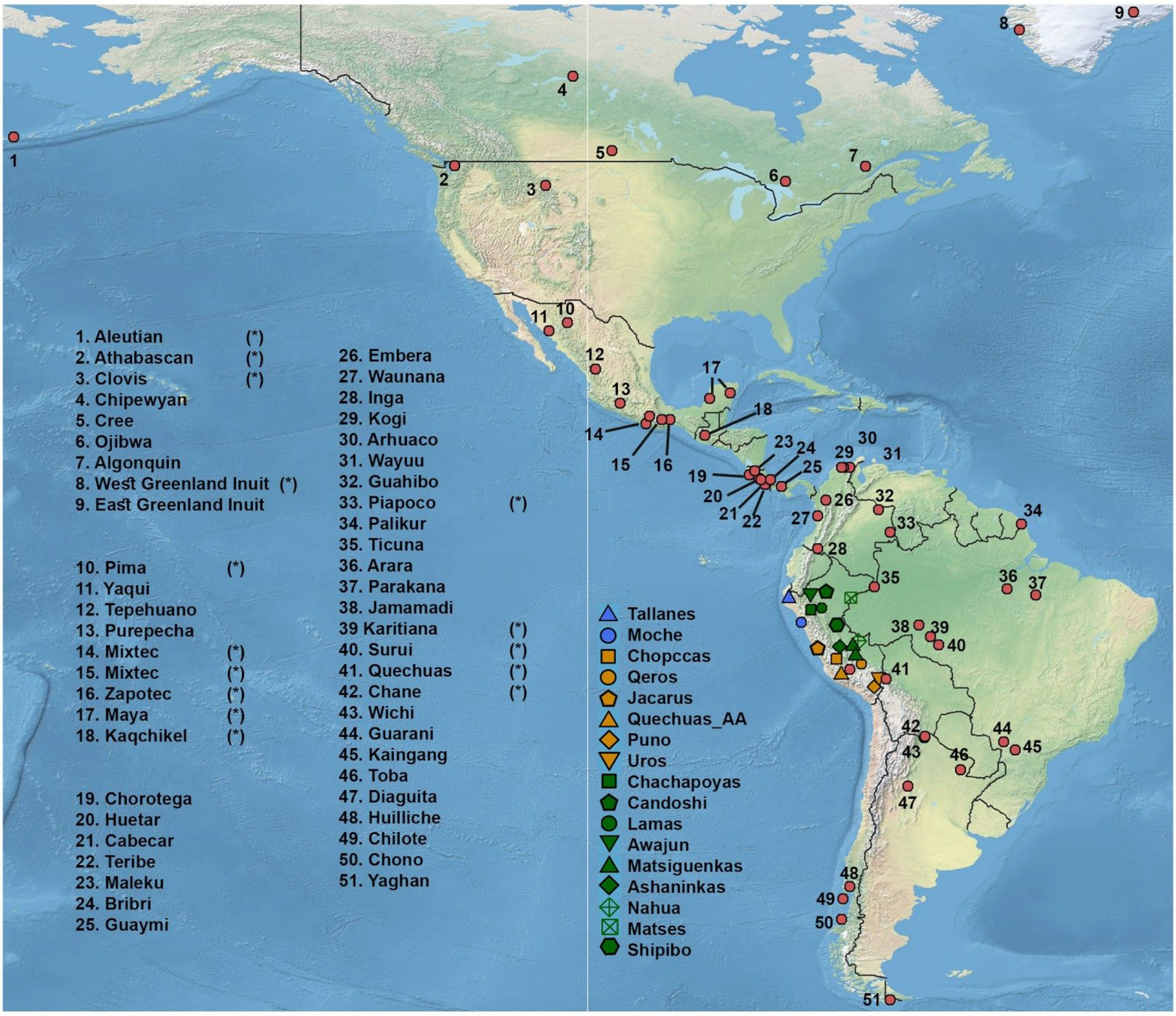


**Figure S1.** Geographical distribution for the 18 Peruvian Native populations sampled, plus the 65 sampled Native American populations and public data sets (Mallick et al. 2016, Raghavan et al. 2015, Reich et al. 2012). All samples except Clovis and Athabascan were included in a data set of ~ 230,000 SNPs. Peruvian samples and (*) were included in a data set of ~ 500,000 SNPs. Matsiguenkas 1= Matsiguenkas-Sepahua; Matsiguenkas 2= Matsiguenkas-Shimaa.

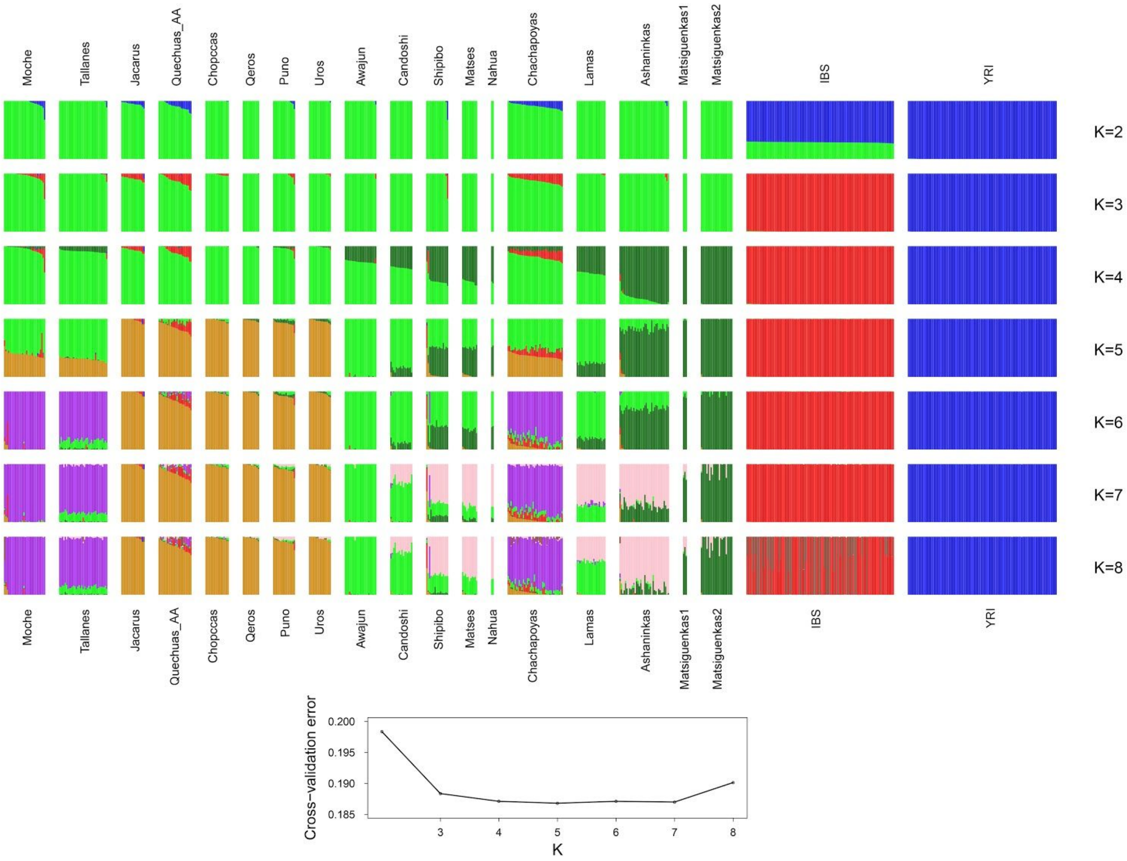


**Figure S2.** ADMIXTURE analysis for 18 Native American populations, as well as Iberian (IBS) and Yoruba (YRI) populations from 1000 Genomes Project (Natives 1.9M Dataset). Figure shows results for 2 to 8 ancestral clusters (K) and a plot (Bottom) with the ADMIXTURE cross-validation errors as a function of K. The lowest cross validations error corresponds to K=5 in which we observed four Native American, one European and one African cluster.

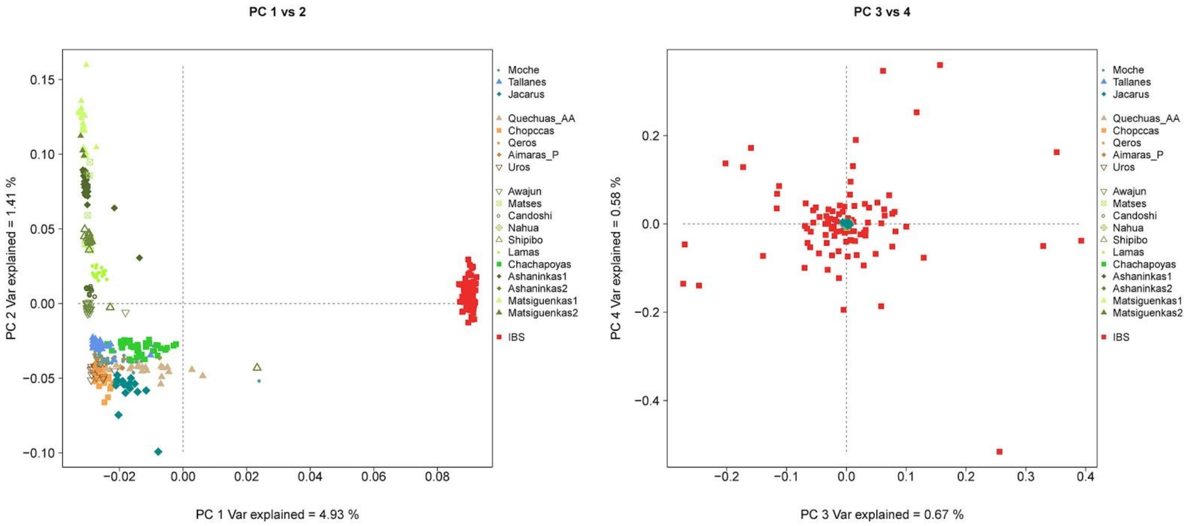


**Figure S3.** Principal Component Analysis for 18 Native American Peruvian populations and Iberian individuals (IBS) from 1000 Genomes Project (Natives 1.9M Dataset). Shades of blue are related to Coast populations. Orange-brown colors are related to Andean populations and green colors are related to Amazon.

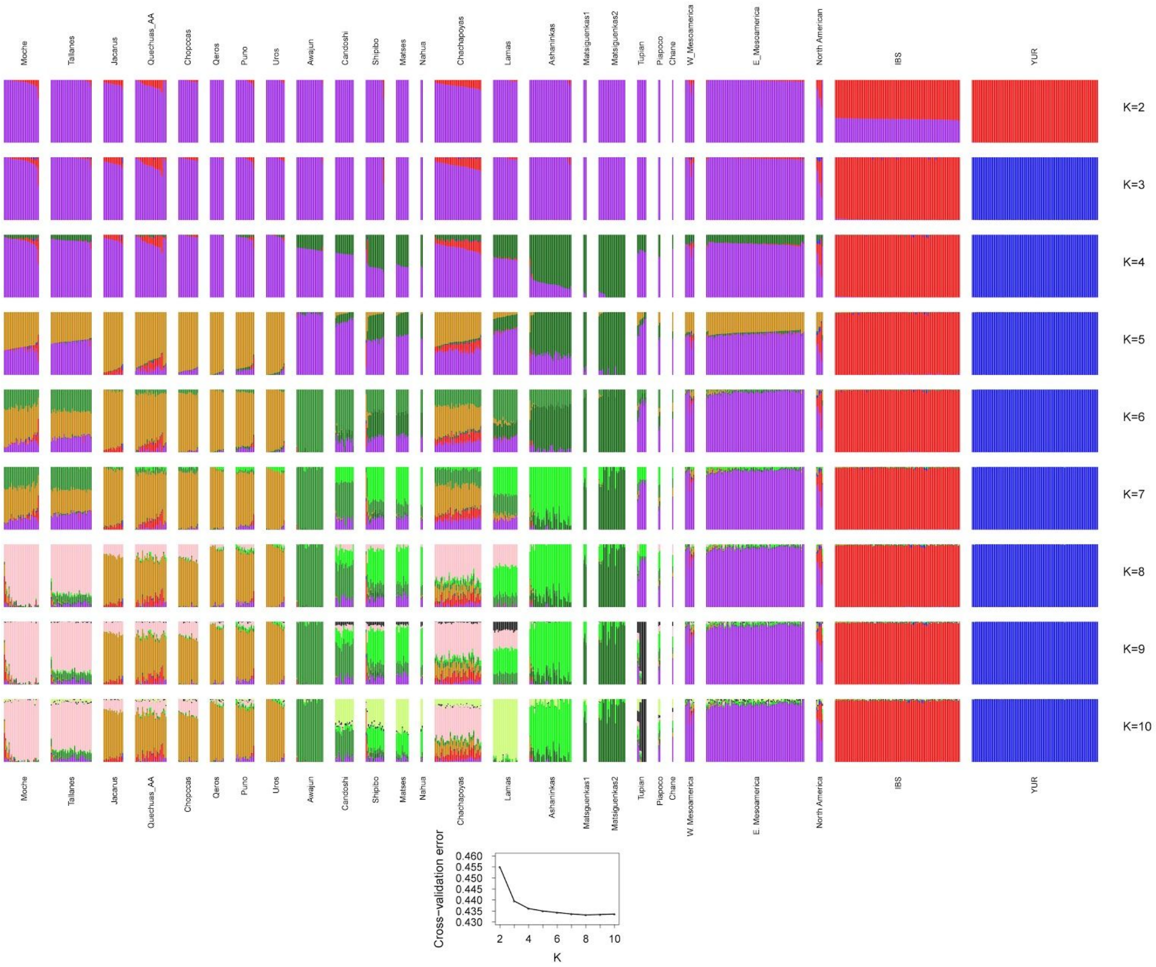


**Figure S4.** ADMIXTURE analysis for 18 Natives American Peruvian populations, Guatemala samples, Native Americans from Raghavan et al. 2015 and the Simons Project (Mallick et al. 2016) Iberian (IBS) and Yoruba (YRI) populations from 1000 Genomes Project (Natives 500K Dataset). Figure shows results for 2 to 10 ancestral (K) clusters and a plot (Bottom) with the ADMIXTURE cross-validation errors as a function of K. The lowest cross validation error corresponds to K=8.

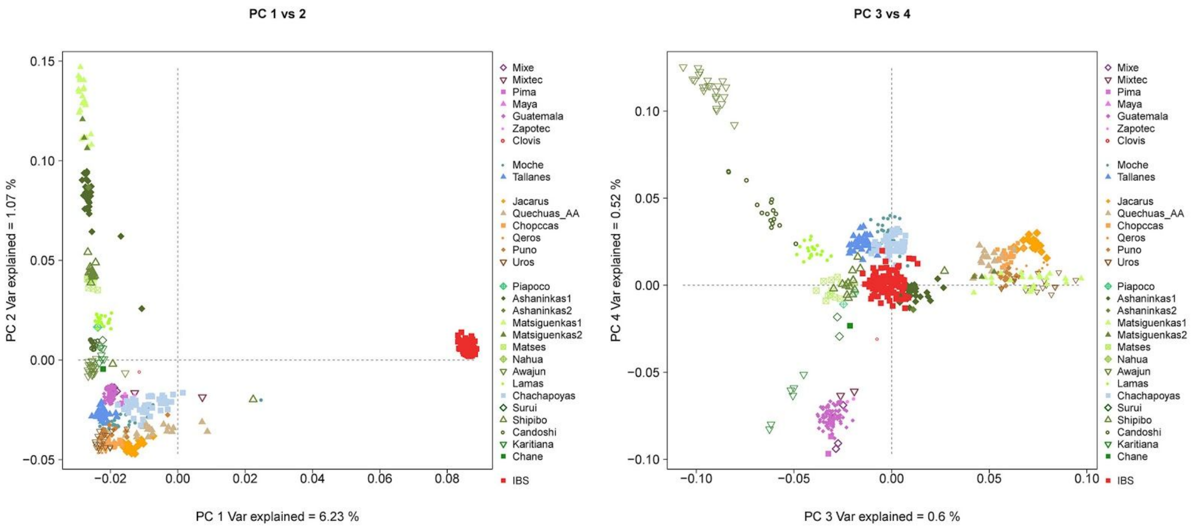


**Figure S5.** Principal Component Analysis for 18 Native American Peruvian populations, Guatemala samples, Native Americans from Raghavan et al. 2015 and the Simons Project (Mallick et al. 2016) and Iberian (IBS) population from 1000 Genomes Project (Natives 500K Dataset). Shades of blue are related to Peruvian Coast populations. Orange-brown colors are related to Andean populations and green colors are related to Amazon. Shades of purple are related to Mesoamericans. Shades of beige are related to North American natives.

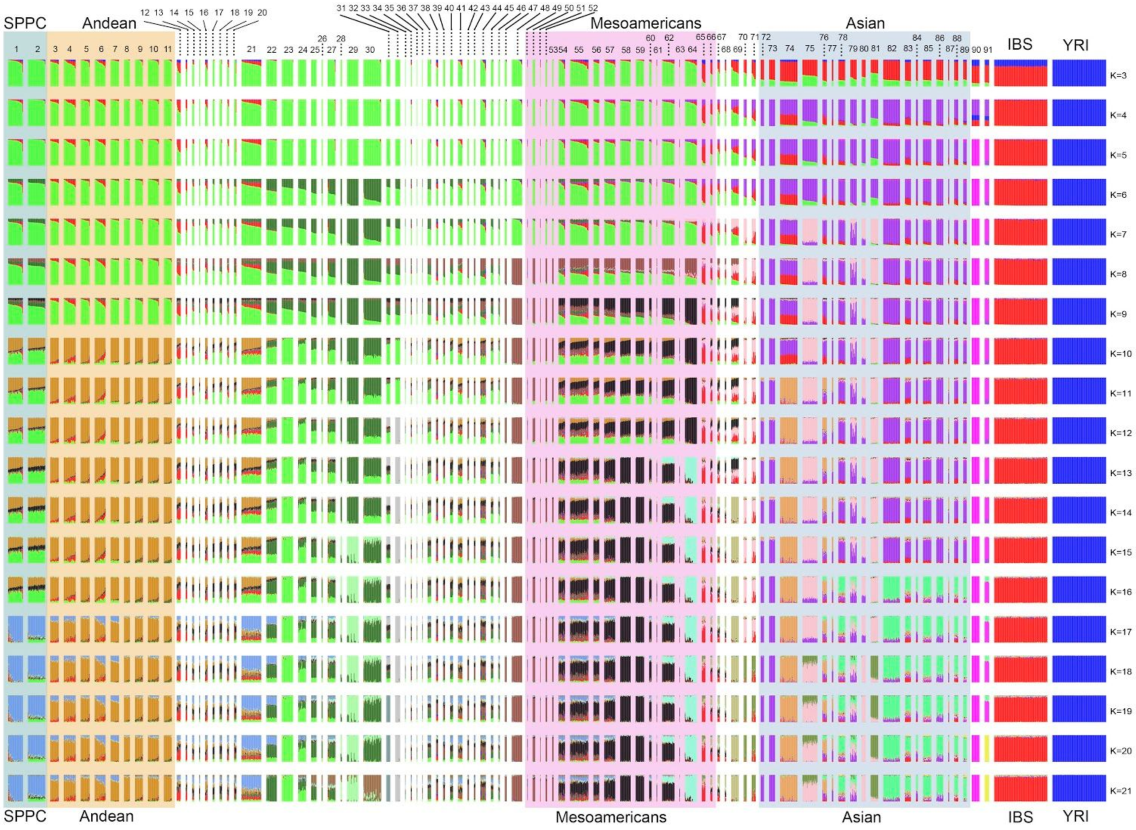


1. Moche
2. Tallanes
3. Jacarus
4. Quechuas
5. Quechua_R1
6. Quechua_R2
7. Chopccas
8. Qeros
9. Aimaras
10. Aimaras_R
11. Uros
12. Chilote
13. Chono
14. Hulliche
15. Chane
16. Diaguita
17. Toba
18. Wichi
19. Yaghan
20. Guarani
21. Chachapoyas
22. Lamas
23. Awajun
24. Candoshi
25. Matses
26. Nahua
27. Shipibo
28. Matsiguenkas
29. Shimaa
30. Ashaninkas
31. Karitiana
32. Surui
33. Arara
34. Kaingang
35. Parakana
36. Jamamadi
37. Piapoco
38. Arhuaco
39. Embera
40. Guahibo
41. Inga
42. Kogi
43. Waunana
44. Wayuu
45. Ticuna
46. Palikur
47. Bribri
48. Cabecar
49. Chorotega
50. Guaymi
51. Huetar
52. Maleku
53. Teribe
54. Kaqchikel
55. Maya1
56. Maya2
57. Zapotec1
58. Zapotec2
59. Mixe
60. Mixtec
61. Purepecha
62. Tepehuano
63. Yaqui
64. Pima
65. Aleutian
66. Cree
67. Ojibwa
68. Algonquin
69. Chipewyan
70. EastGreenland
71. WestGreenland
72. Nganasan1
73. Nganasan2
74. Khanty
75. Chukchi

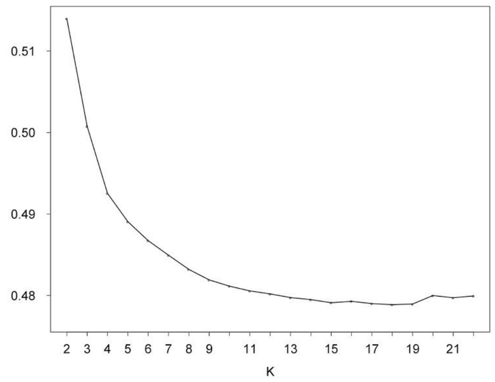


76. Selkup
77. Tundra_Nentsi
78. Tuvinians
79. Yukaghir
80. Koryak
81. Naukan
82. Yakut
83. Altaian
84. Dolgan
85. Buryat
86. Evenki
87. Ket
88. Mongolian
89. Mongola
90. Papuan
91. Melanesian
92. IBS
93. YRI

**Figure S6.** (Top) ADMIXTURE analysis for 90 worldwide populations including 71 Native American populations, 18 Asian, 2 Oceanian, Iberian (IBS) and Yoruba (YRI) populations from 1000 Genomes Project (Natives 230K Dataset). Figure shows results for 3 to 21 ancestral clusters (K). (Bottom) ADMIXTURE cross-validation errors as a function of K and list of populations included. The lowest cross validations corresponds to ADMIXTURE K=18.

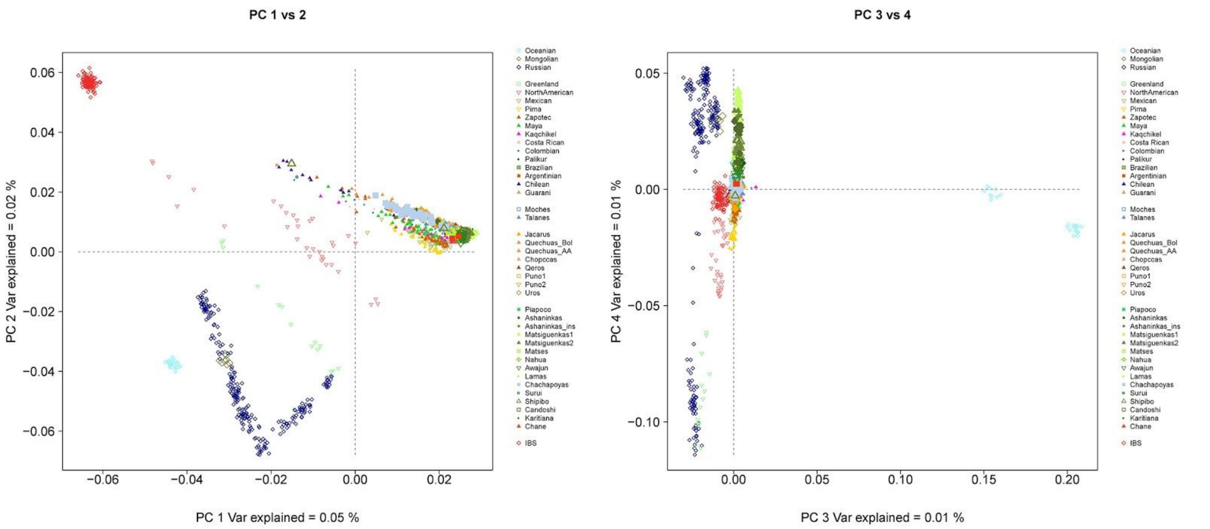


**Figure S7.** Principal Component Analysis for 89 worldwide populations including 68 Native American populations, 18 Asian populations, 2 Oceanian populations, Iberian (IBS) populations (Native 230K Dataset).

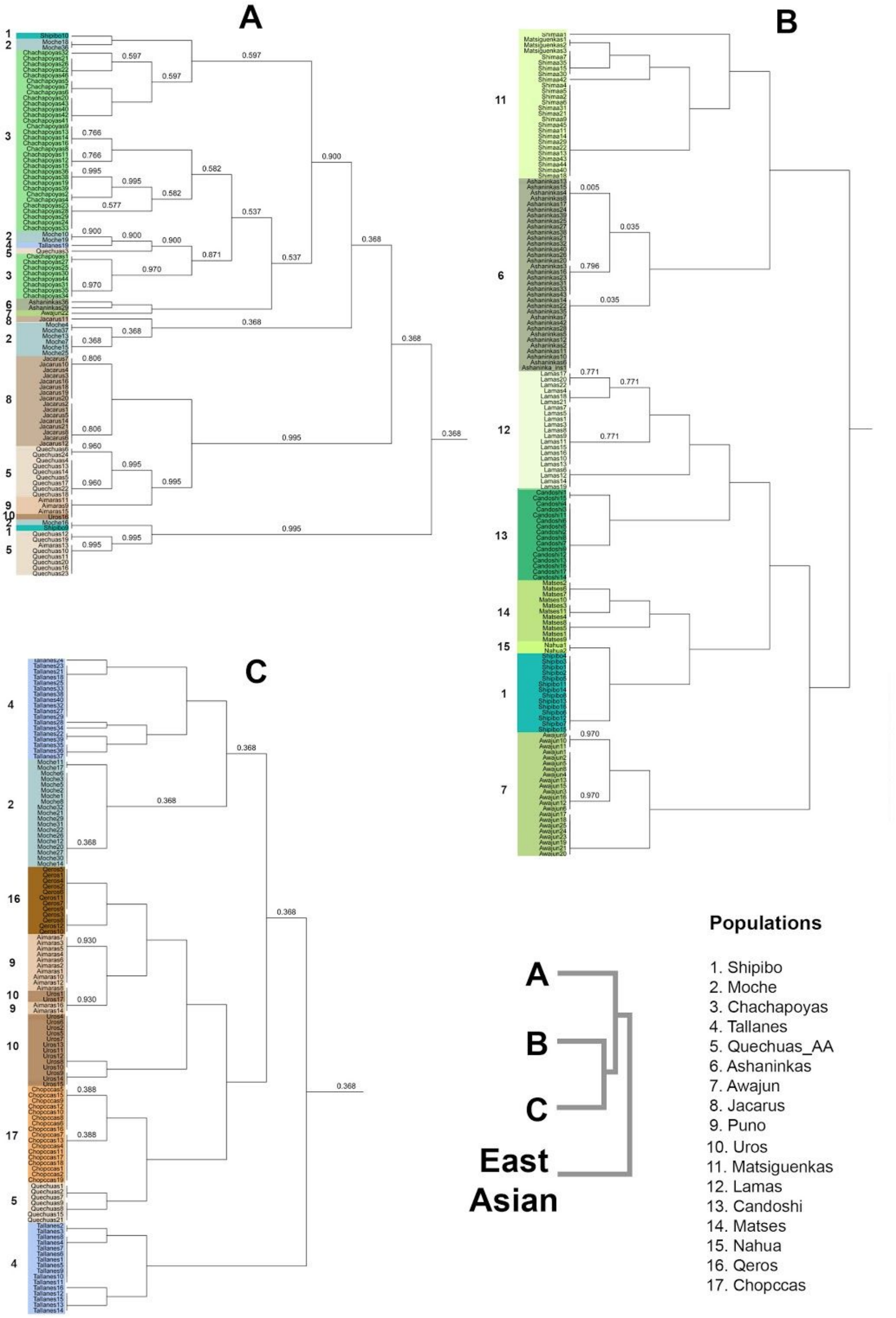


**Figure S8.** fineSTRUCTURE clustering analysis for the Dataset 1.9M dataset. The tree shows the haplotype sharing between Native Americans and East Asian samples. Figures A, B, and C represent the clusters A, B, and C, respectively, in the tree on the right. East Asian clusters grouped all Asian samples. Shades of blue are related to Peruvian Coast populations. Orange-brown colors are related to Andean populations and green colors are related to Amazon. Shades of purple are related to Mesoamericans. Shades of beige are related to North American natives.

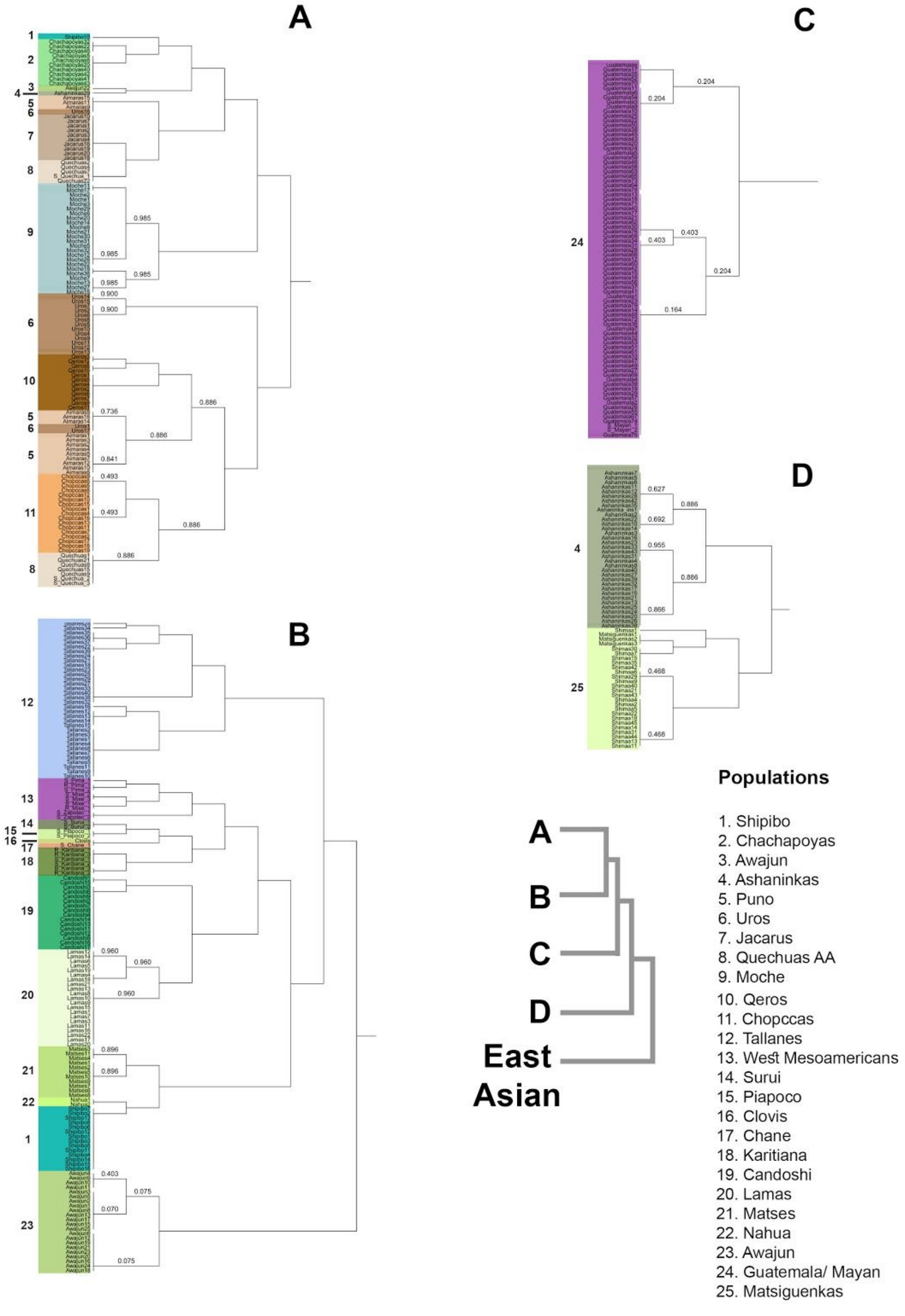


**Figure S9.** fineSTRUCTURE clustering analysis for the Dataset 500K dataset. The tree shows the haplotype sharing between Native Americans and East Asian samples. Figures A, B, C and D represent the clusters A, B, C and D, respectively, in the tree on the right. East Asian clusters grouped all Asian samples. Shades of blue are related to Peruvian Coast populations. Orange-brown colors are related to Andean populations and green colors are related to Amazon. Shades of purple are related to Mesoamericans. Shades of beige are related to North American natives.

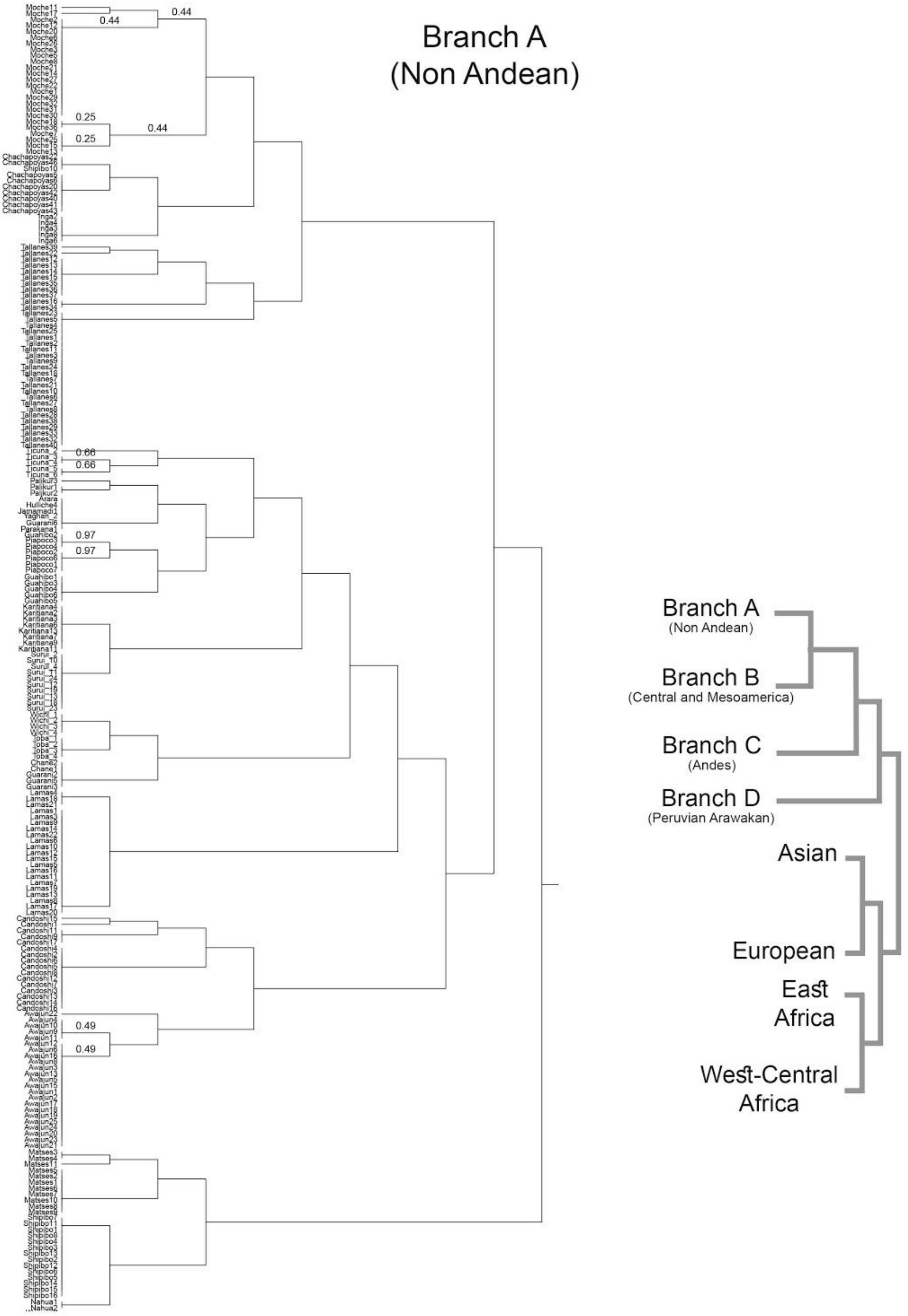


**Figure S10.** fineSTRUCTURE clustering analysis for the Dataset 230K dataset. A) Branch A of the tree showing the clustering of the Non Andean populations of South America. B) Branches B (Central Americans and Mesoamericas), C (Andean populations) and D (Peruvian Arawakan Ashaninkas and Matsiguenkas).

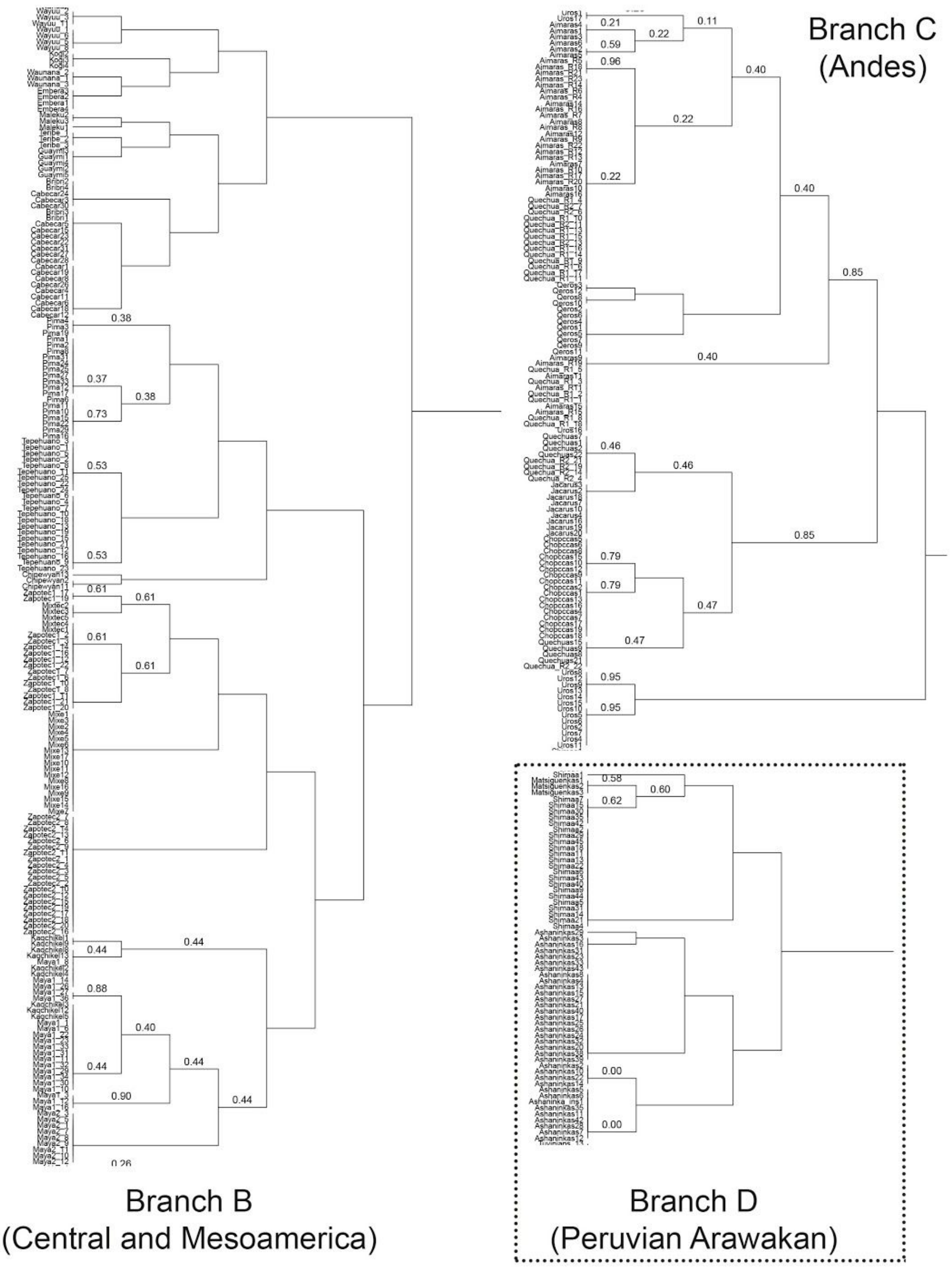


**Figure S10 (Continued).** fineSTRUCTURE clustering analysis for the Dataset 230K dataset. A) Branch A of the tree showing the clustering of the Non Andean populations of South America. B) Branches B (Central Americans and Mesoamericas), C (Andean populations) and D (Peruvian Arawakan Ashaninkas and Matsiguenkas).

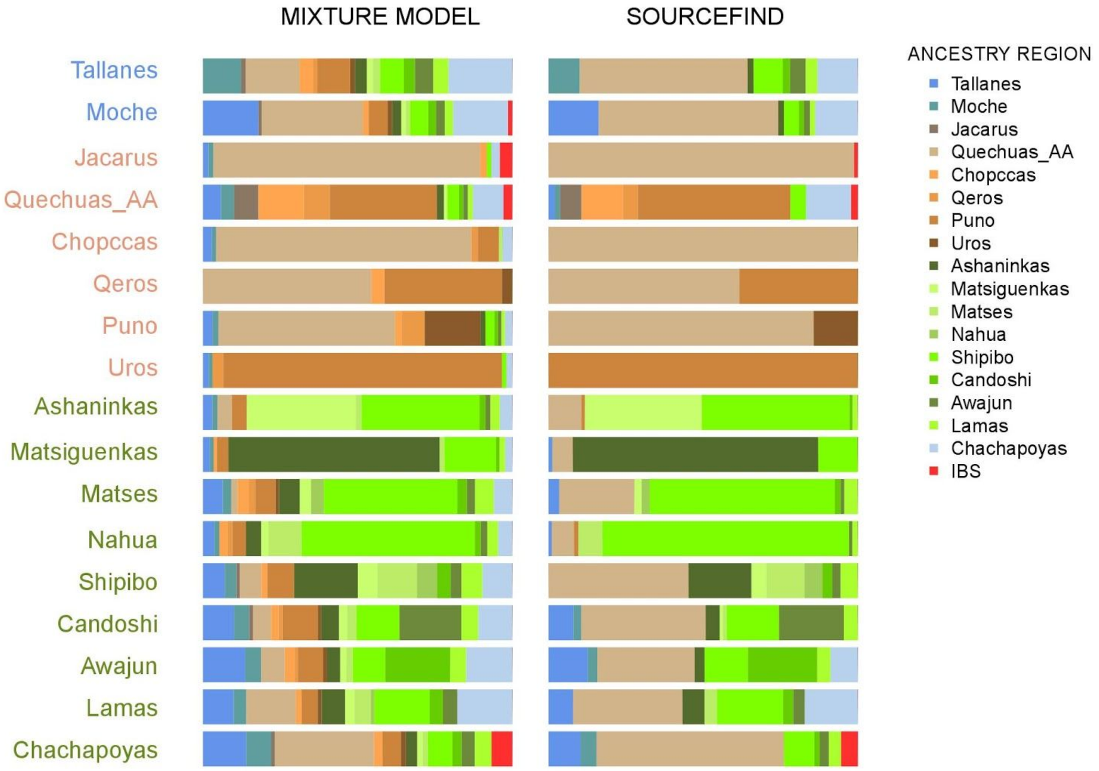


**Figure S11.** Proportions of haplotype sharing for each target population respect to Native Americans, Europeans and Africans donors populations, for the Dataset Natives 1.9M, inferred by two approaches: A non negative regression (MIXTURE MODEL) and a Bayesian approach (SOURCEFIND). Colored bars indicate a proportion of shared haplotypes shared DNA between the target population and a specific donor.

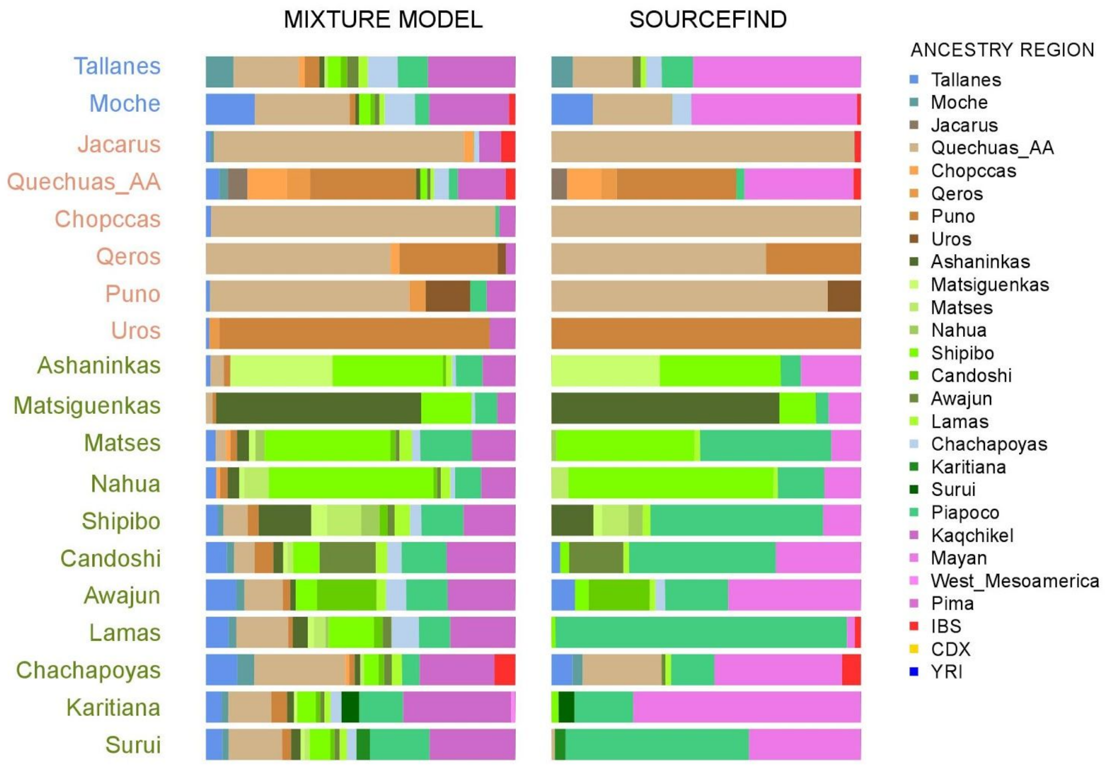


**Figure S12.** Proportions of haplotype sharing for each target population respect to Native Americans, Europeans and Africans donors populations, for the Dataset Natives 500K, inferred by two approaches: A non negative regression (MIXTURE MODEL) and a Bayesian approach (SOURCEFIND). Colored bars indicate a proportion of shared haplotypes shared DNA between the target population and a specific donor.

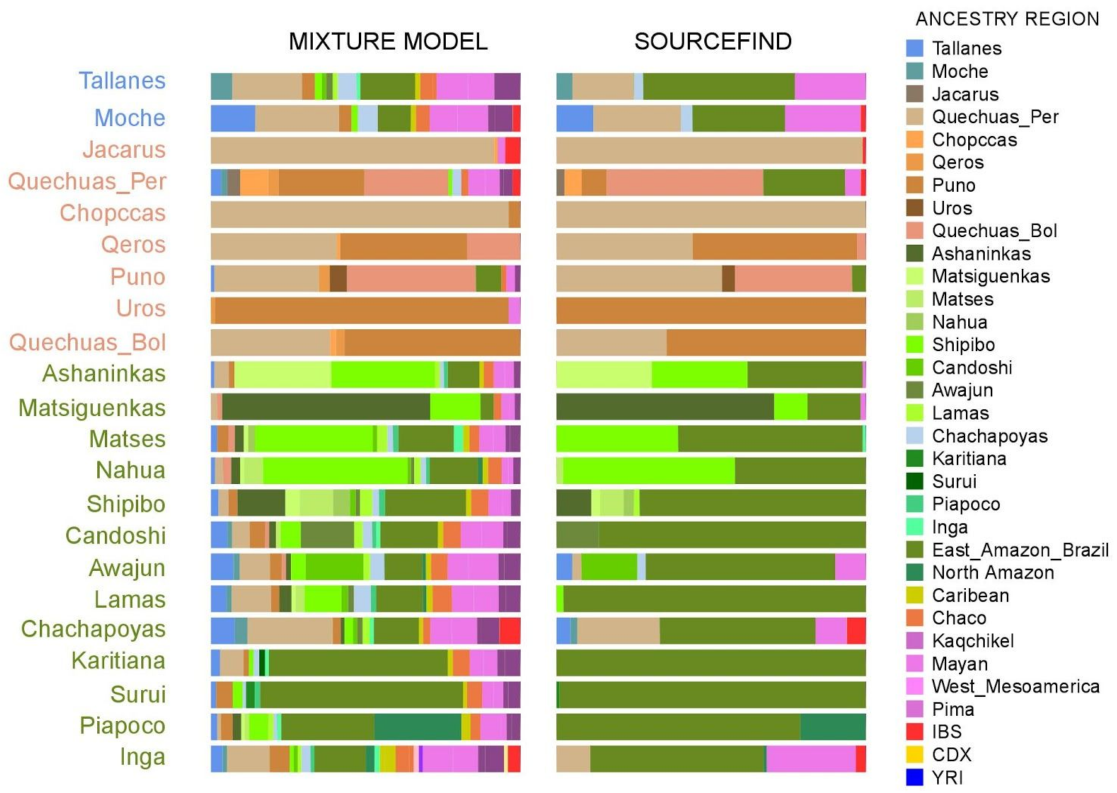


**Figure S13.** Proportions of haplotype sharing for each target population respect to Native Americans, Europeans and Africans donors populations, for the Dataset Natives 230K, inferred by two approaches: A non negative regression (MIXTURE MODEL) and a Bayesian approach (SOURCEFIND). Colored bars indicate a proportion of shared haplotypes shared DNA between the target population and a specific donor.

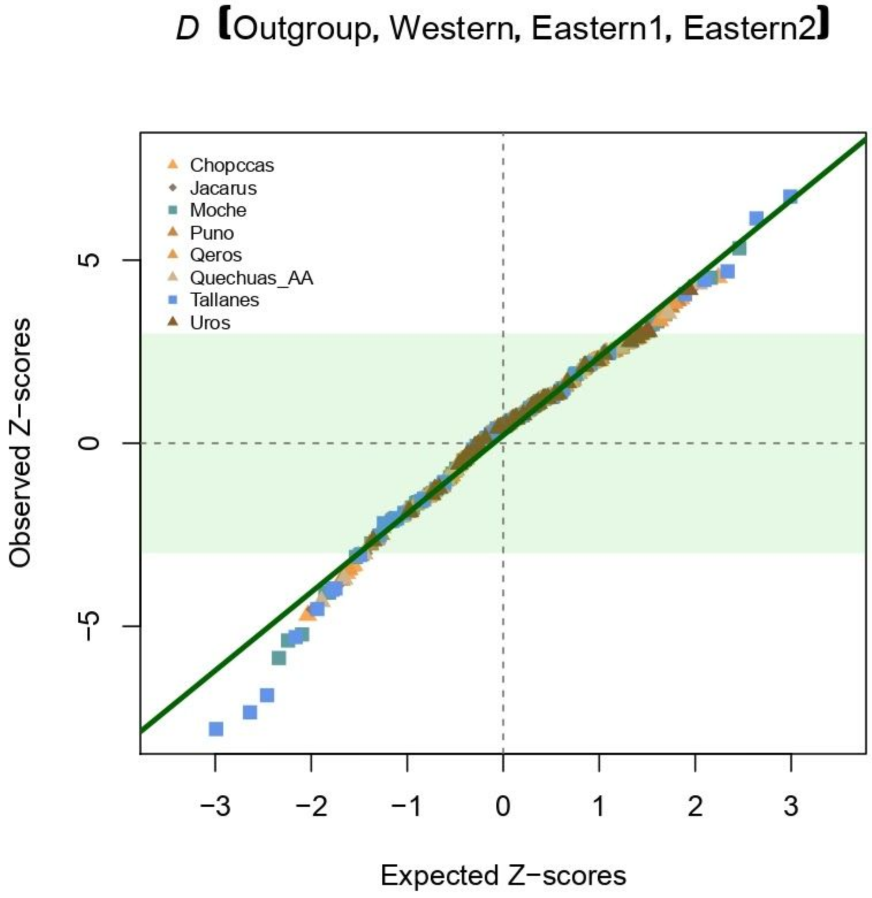


**Figure S14.** Quantile-quantile plot comparing Z-scores from D-statistics relating Western and Eastern Andean slope populations to those expected under a normal distribution for the Dataset 500K. We tested the configuration (outgroups (Western (Eastern1, Eastern2))). We detected strong genetic affinity between Peruvian Coast and Eastern populations in the North Fertile Andes.

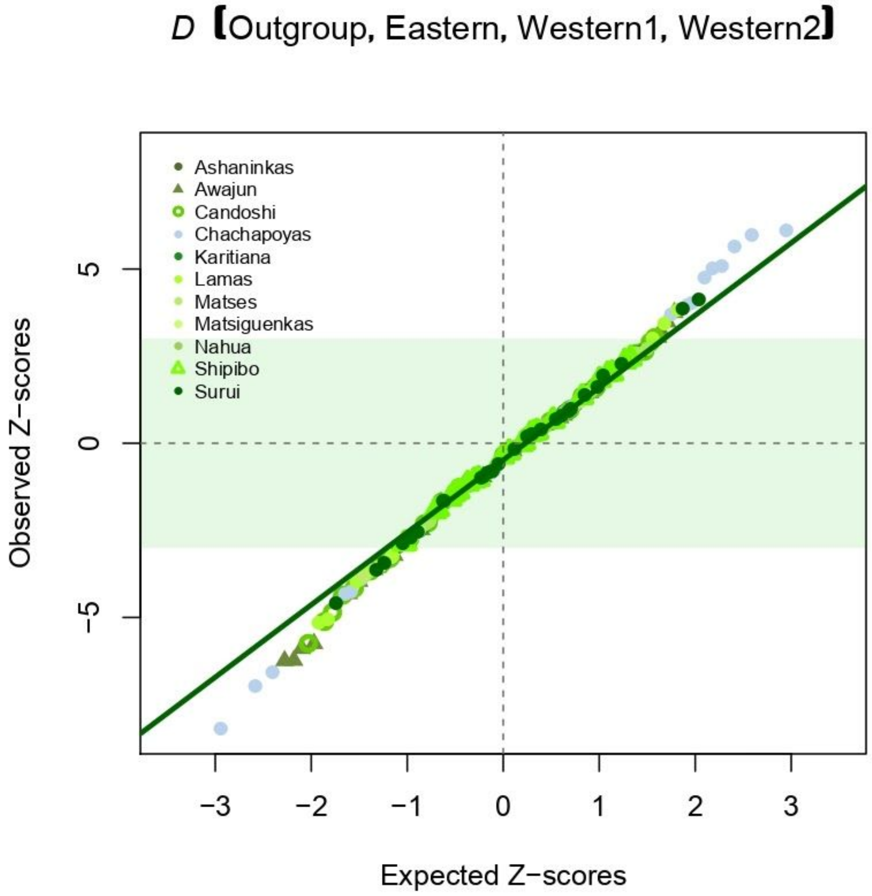


**Figure S15:** Quantile-quantile plot comparing Z-scores from D-statistics relating Western and Eastern Andean slope populations to those expected under a normal distribution for the Dataset 500K. We test the configuration (outgroups (Eastern (Western1, Western2))). We detected strong genetic affinity between Awajun, Candoshi, Lamas and Chachapoyas with Western populations.

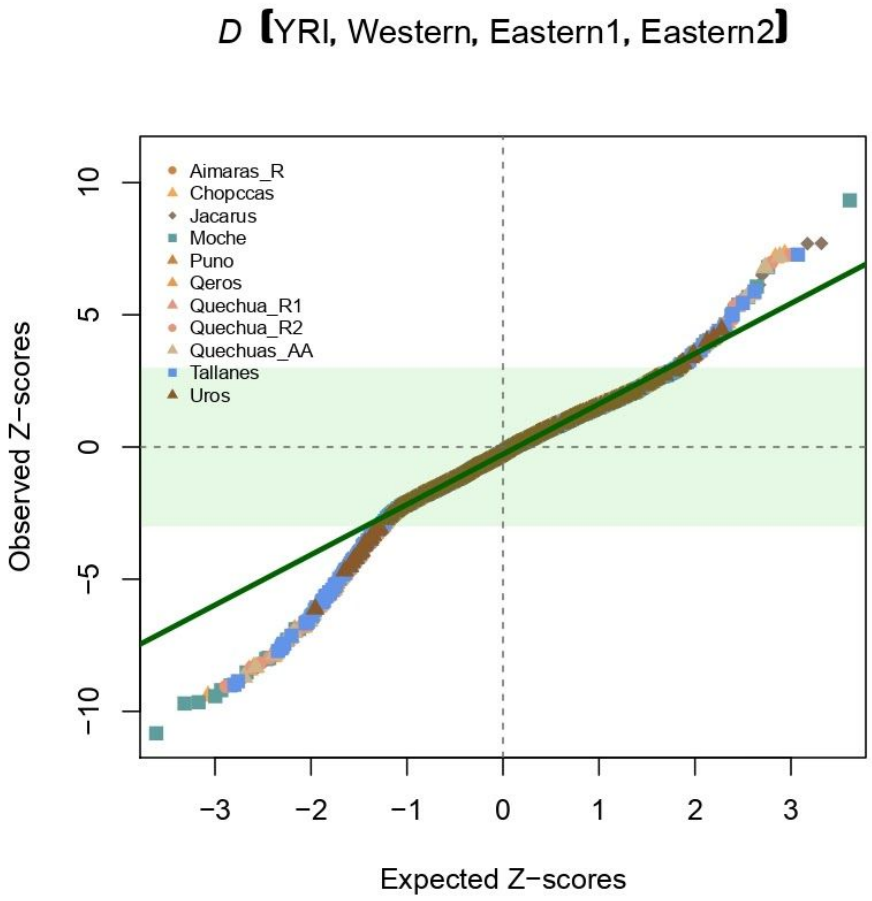


**Figure S16:** Quantile-quantile plot comparing Z-scores from D-statistics relating Western and Eastern Andean slope populations to those expected under a normal distribution for the Dataset 230K. We test the configuration (outgroups (Western (Eastern1, Eastern2))). We detected strong genetic affinity between Peruvian Coast and Eastern populations in the North Fertile Andes.

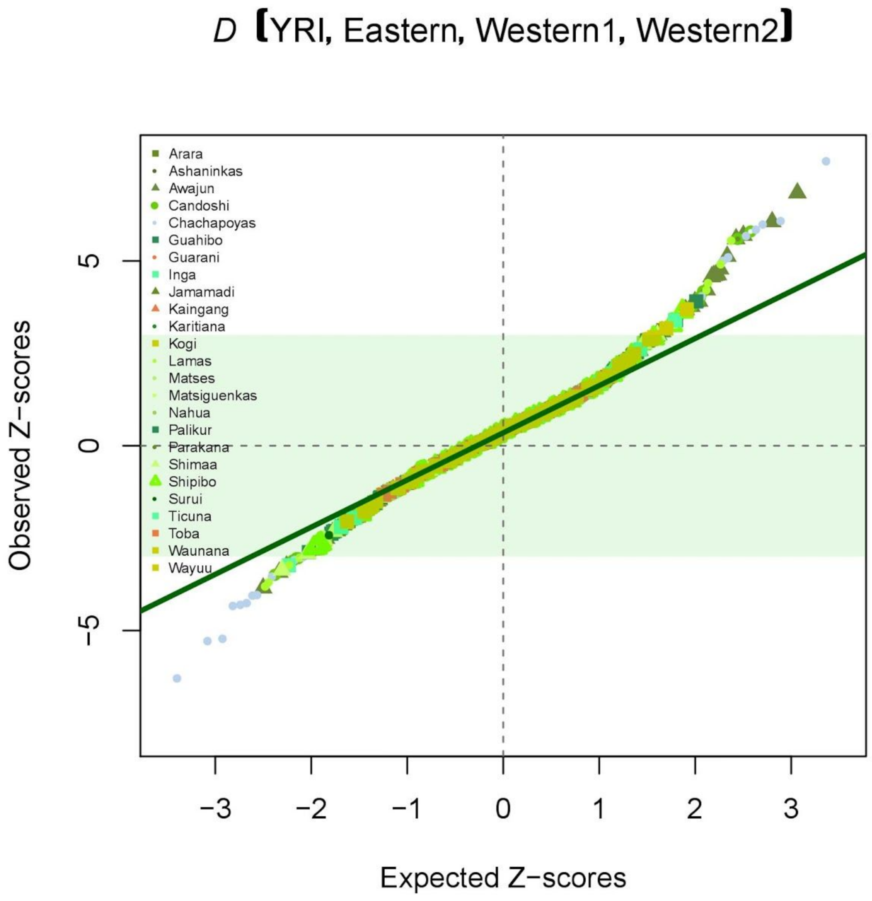


**Figure S17:** Quantile-quantile plot comparing Z-scores from D-statistics relating Western and Eastern Andean slope populations to those expected under a normal distribution for the Dataset 230K. We test the configuration (outgroups (Eastern (Western1, Western2))). We detected strong genetic affinity between Awajun, Candoshi, Lamas and Chachapoyas with Western populations.

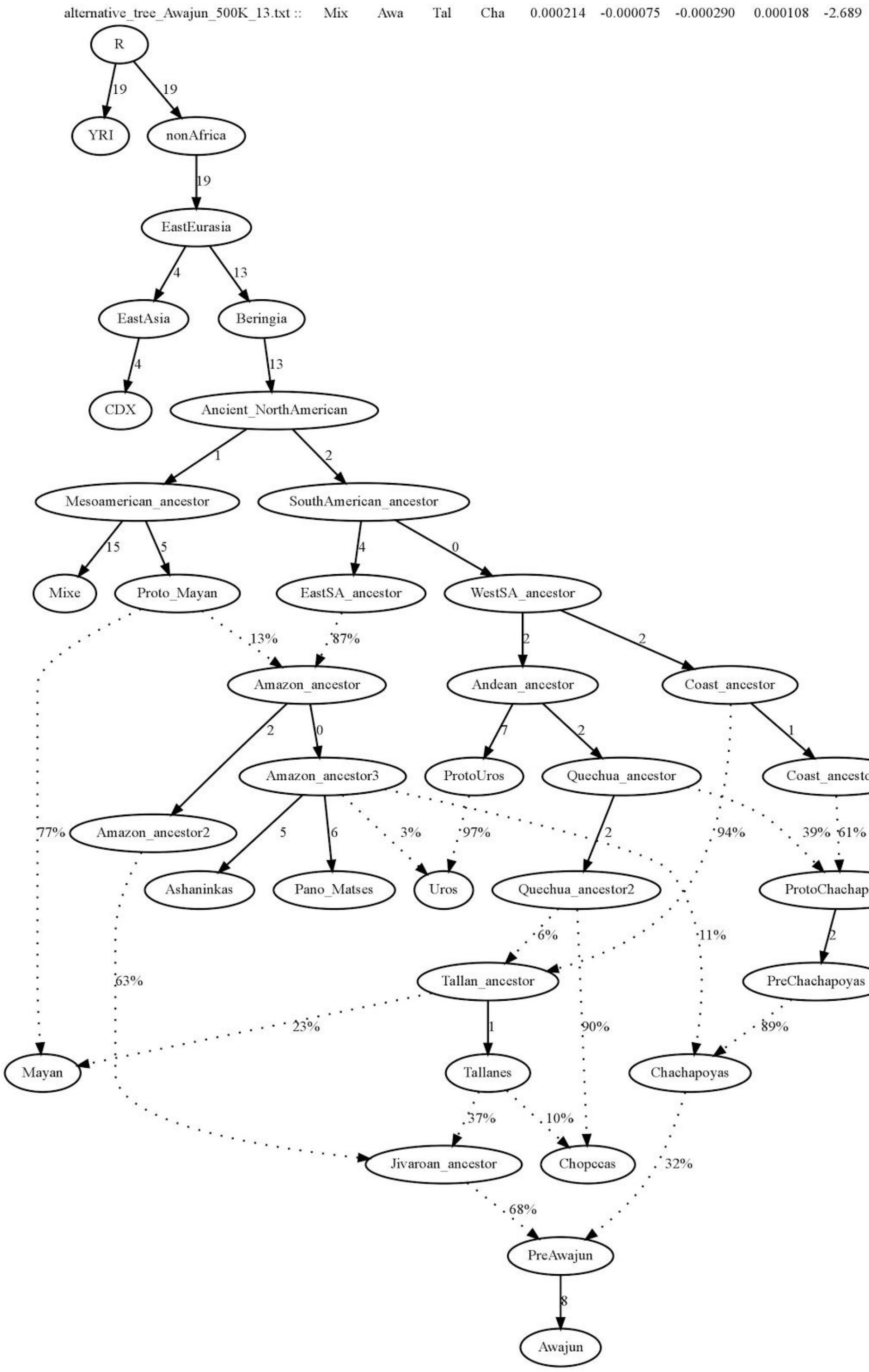


**Figure S18: Admixture Graph model that fits Awajun and Chachapoyas populations in a South American context.** This admixture graph (Z= -2.689) inferred for the Dataset 500K showed the complex relationships of gene flow among Western and Eastern Slope populations. It is observed that Awajun population have an admixed nature from both slopes of the Andes.

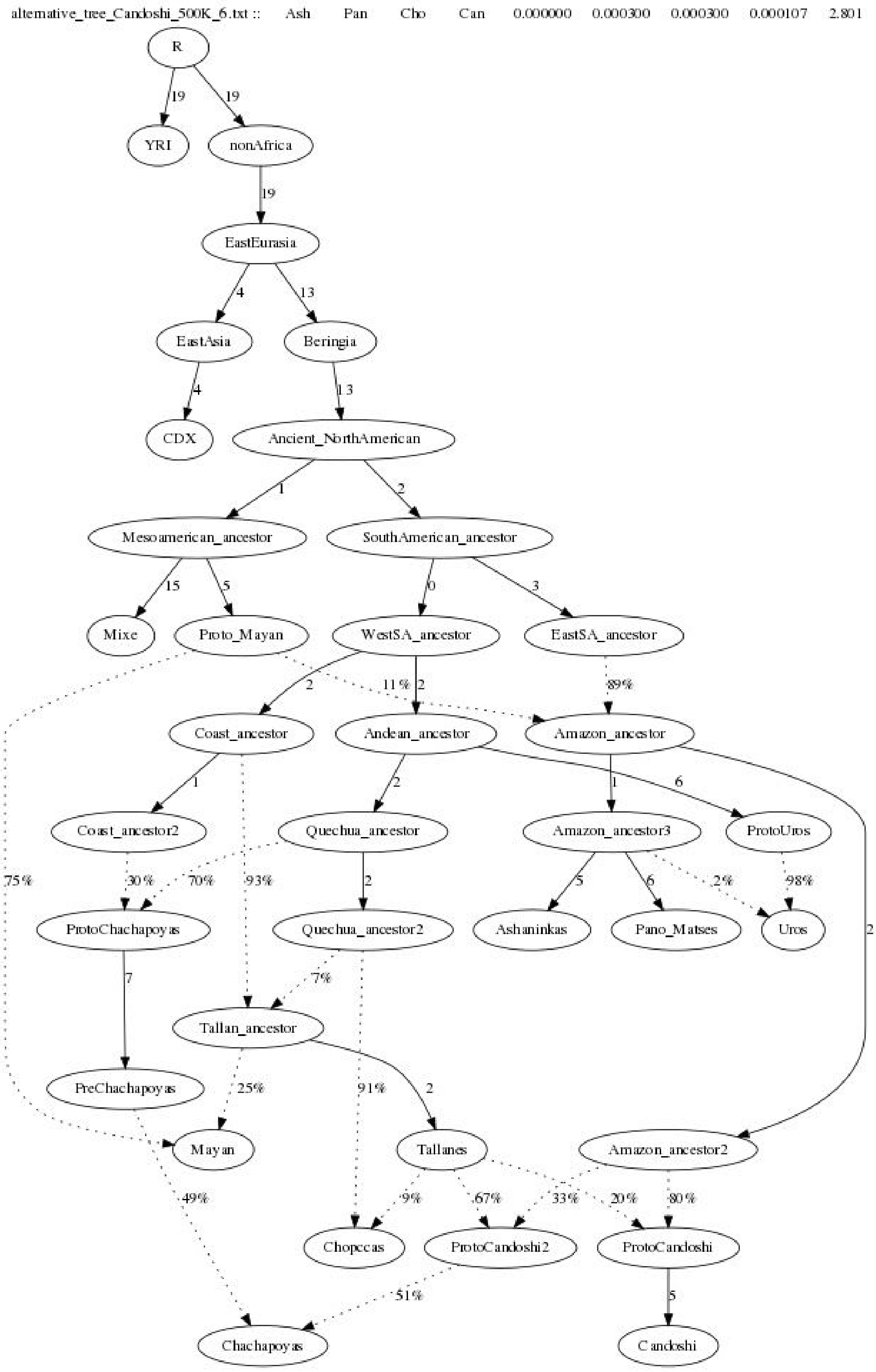


**Figure S19: Admixture Graph model that fits Candoshi and Chachapoyas populations in a South American context.** This admixture graph (Z= 2.801) inferred for the Dataset 500K showed the complex relationships of gene flow among Western and Eastern Slope populations. It is observed that Candoshi population have an admixed nature from both slopes of the Andes.

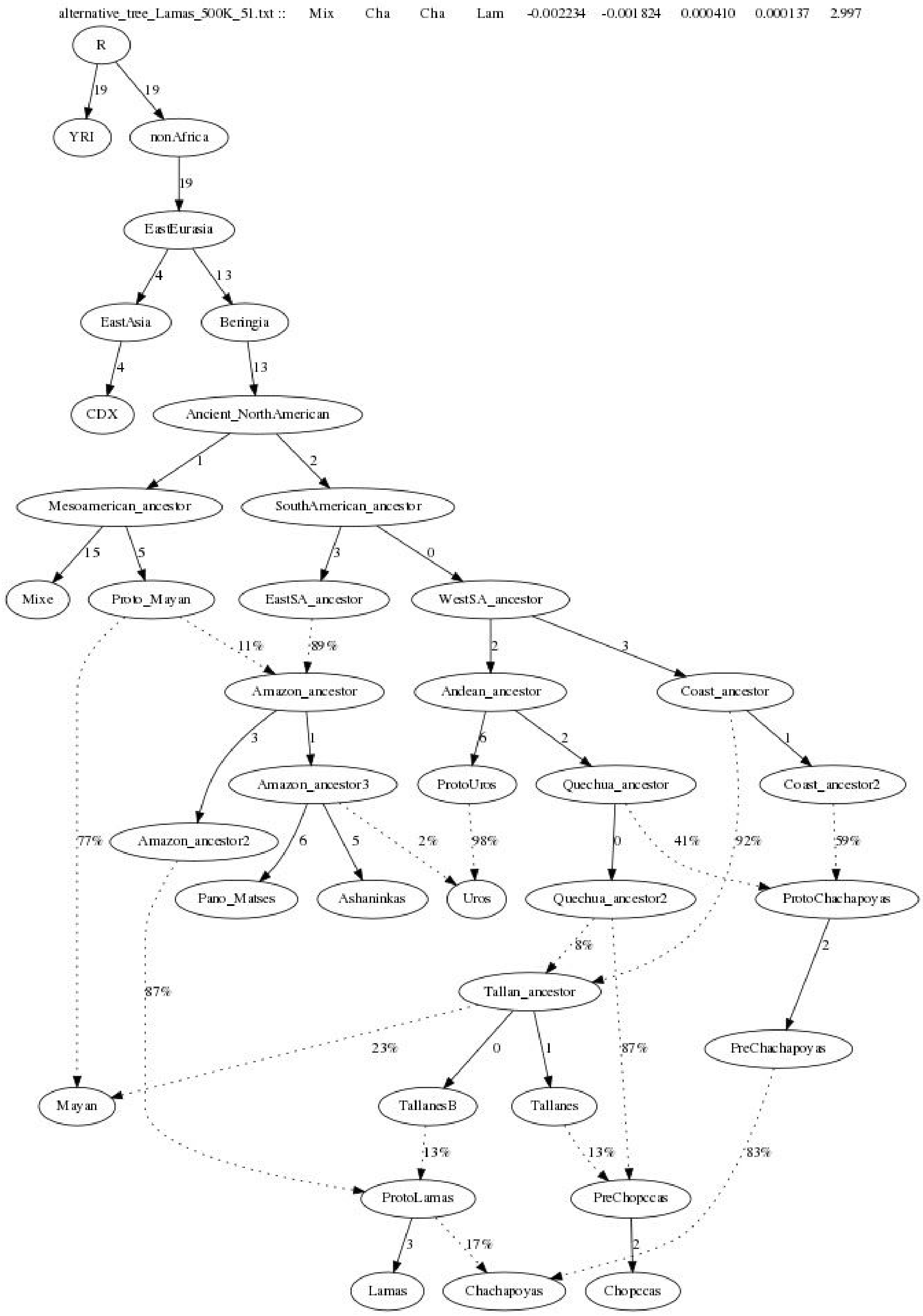


**Figure S20: Admixture Graph model that fits Lamas and Chachapoyas populations in a South American context.** This admixture graph (Z= 2.997) inferred for the Dataset 500K showed the complex relationships of gene flow among Western and Eastern Slope populations. It is observed that Lamas population have an admixed nature from both slopes of the Andes.

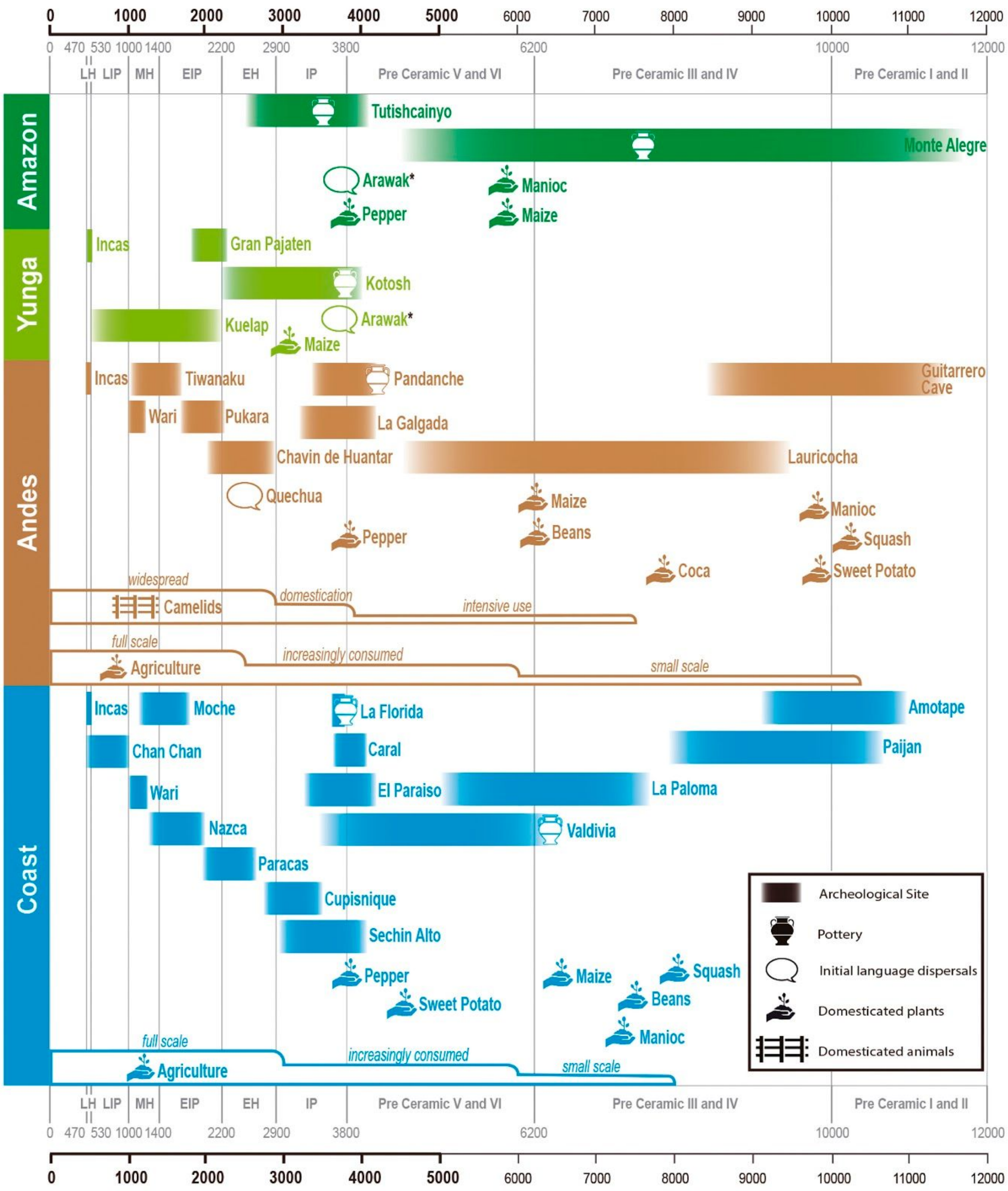


**Figure S21: Key historical events of Peruvian prehistory in four longitudinal regions: Peruvian Coast, Andes, Amazon Yunga and Amazonia.** Pottery and cultivars symbols represent the earliest archaeological record for the region. To account for time uncertainties, This figure showed the events in the chronology plot without clearly defined chronological borders. LH: Late Horizon, LIP: Late Intermediate Period, MH: Middle Horizon, EIP: Early Intermediate Period, EH: Early Horizon, IP: Initial Period. *Controversial geographic region of Arawak origin. Each step in Agriculture and Camelids representations shows an increase in their relative importance. Figure adapted from Scliar et al. 2014.

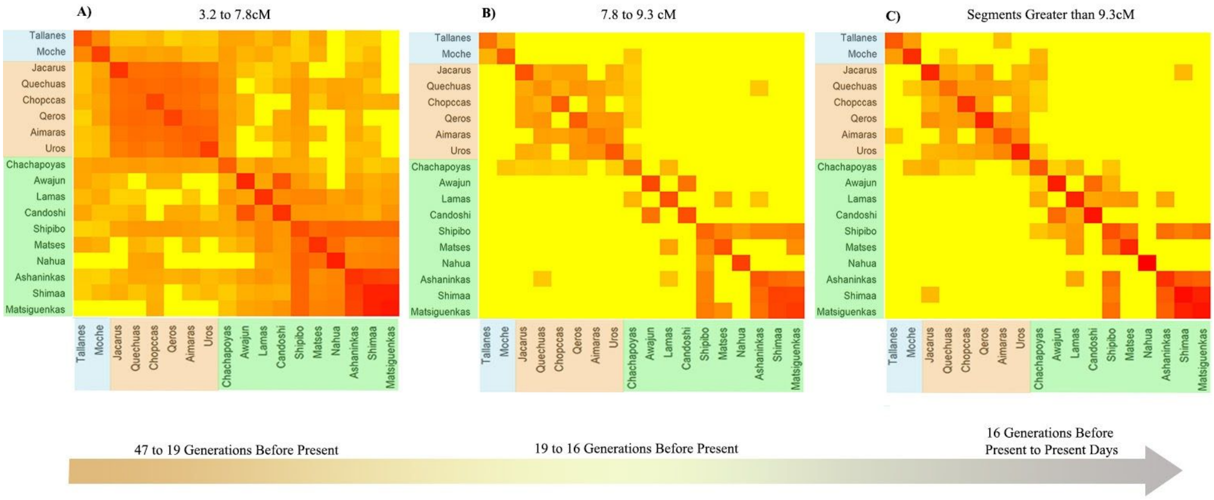


**Figure S22:Heatmap representation of the shared Identical by descent (IBD) segments among Native Americans of the Natives 1.9M dataset.** Each heatmap represents an interval of segments size and is correlated with time generation for the most common recent ancestor. A) An interval from 3.2 to 7.8 cM correlated with 47 to 19 generations ago. B) The second interval from 7.8 to 9.3 cM correlated with 19 to 16 generations ago. C) And the last interval for all segments longer that 9.3 cM correlated with 16 generations ago to the present day.

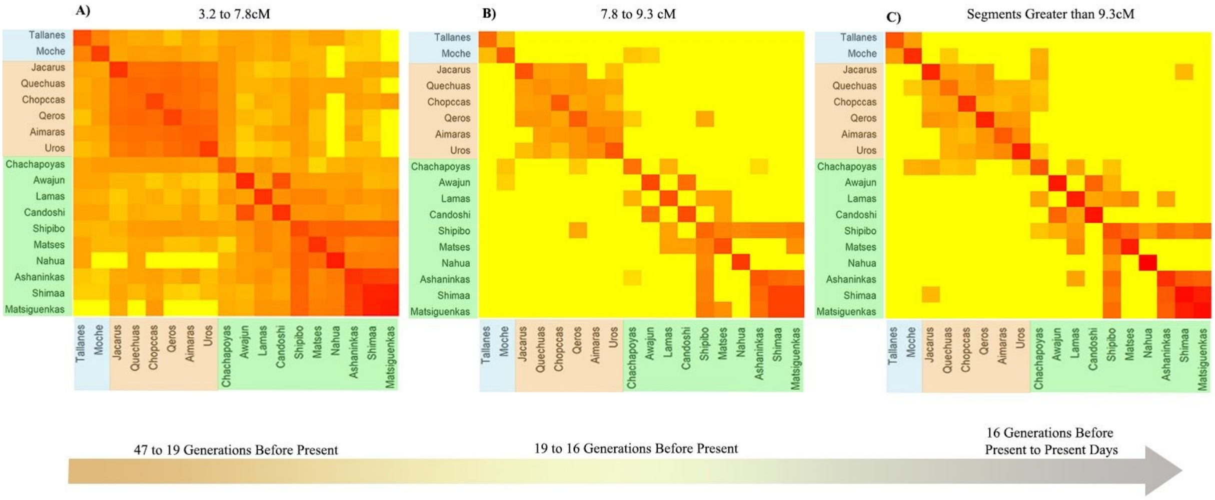


**Figure S23: Heatmap representation of the shared Identical by descent (IBD) segments among Native Americans of the Natives 500K dataset.** Each heatmap represents an interval of segments size and is correlated with time generation for the most common recent ancestor. A) An interval from 3.2 to 7.8 cM correlated with 47 to 19 generations ago. B) The second interval from 7.8 to 9.3 cM correlated with 19 to 16 generations ago. C) And the last interval for all segments longer that 9.3 cM correlated with 16 generations ago to the present day.

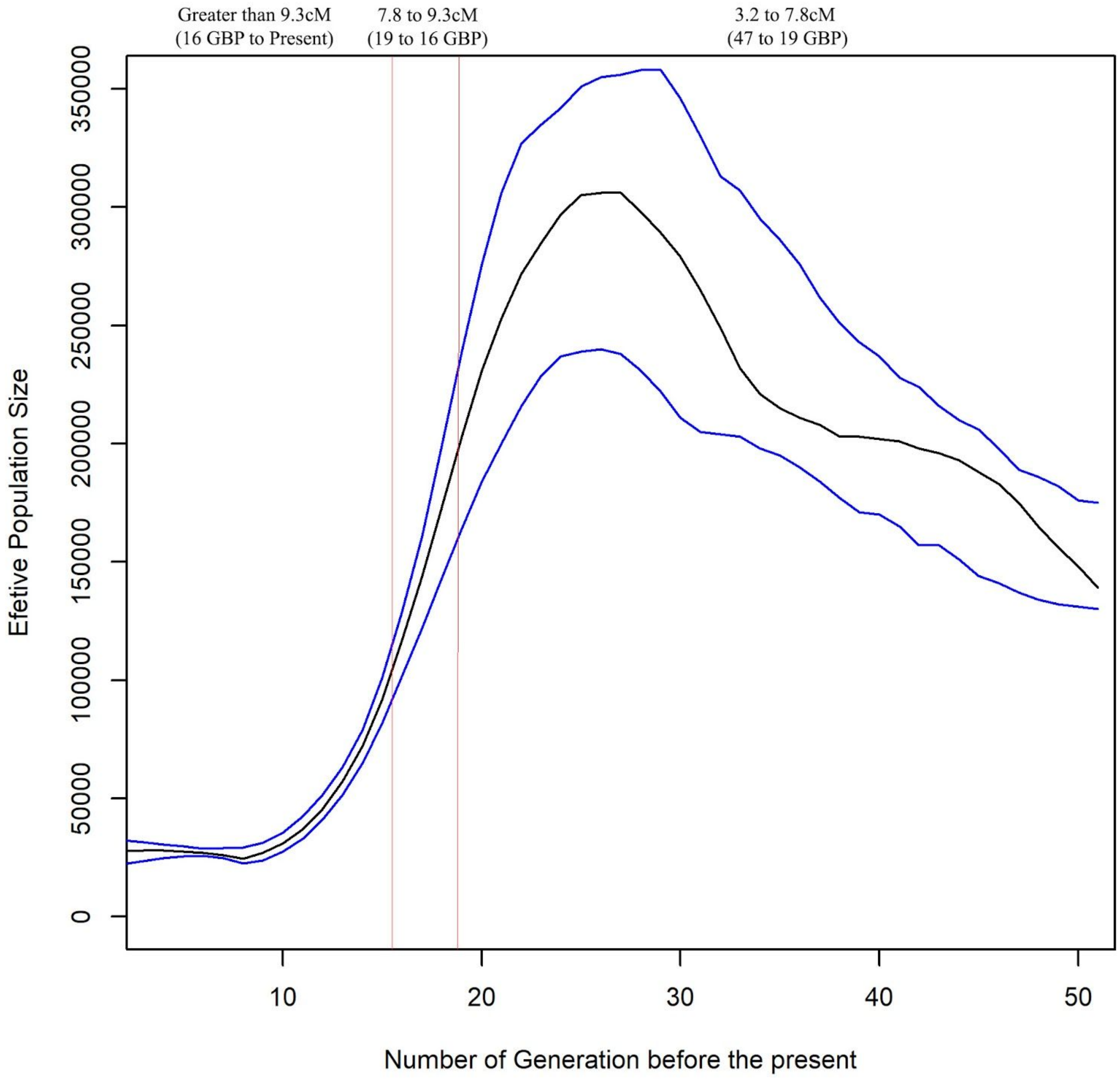


**Figure S24: IBDNe analysis to infer the dynamic of the effective population size (Ne) from 4 generations ago to the last 50 generations for the Andean populations** (Quechuas_AA, Aimaras_P, Chopccas, Qeros and Uros) as a whole. We use the Natives 1.9M dataset. The x axis represents the number of generations from the present to the past. The y axis represents the estimated value of the Ne. Blocks separated by red lines in the graph correspond to the intervals of the IBD heatmaps. GBP: Generations before present.

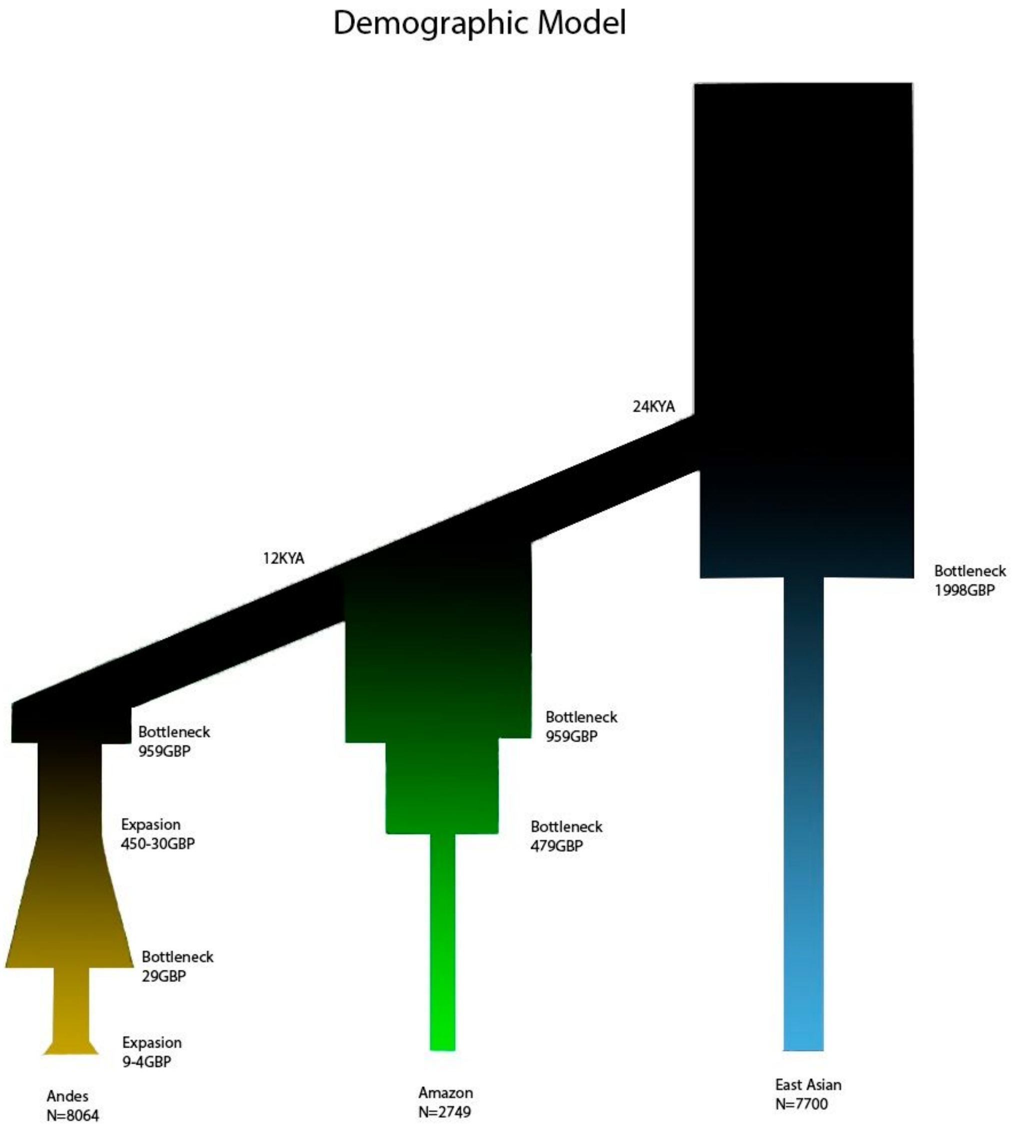


**Figure S25: Demographic model of the Andean, Amazonian and East Asian populations.** This model was used for the simulations made to calculate the p-value of the obtained PBS values.

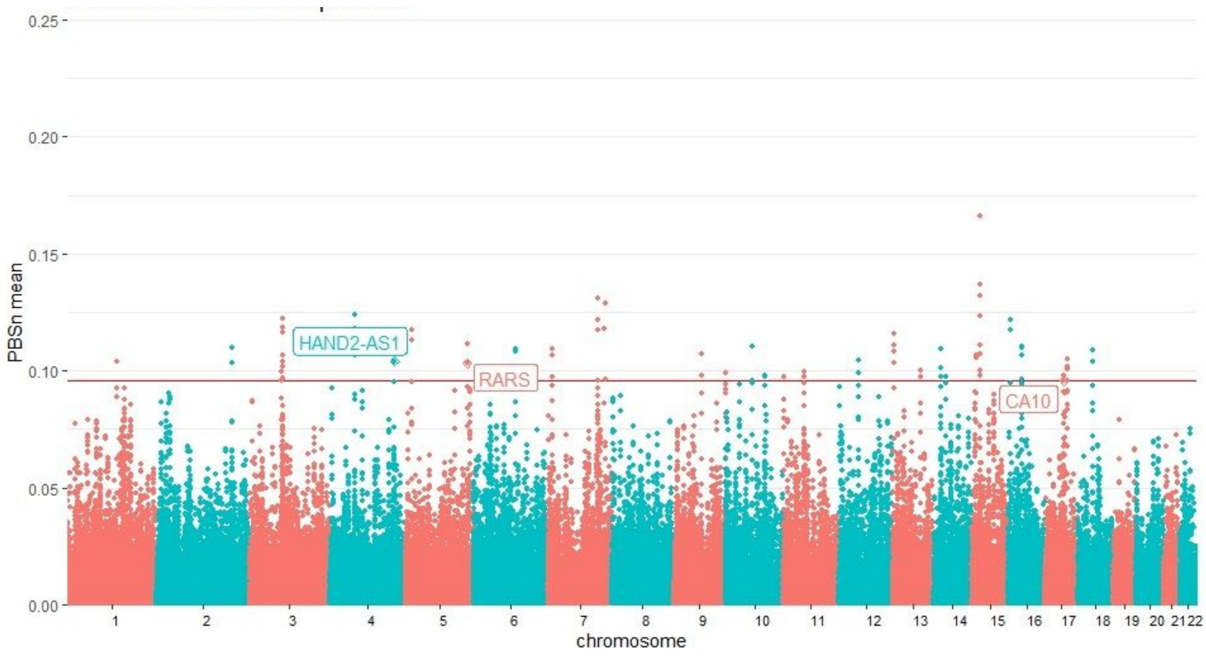


**Figure S26: PBSn mean values for windows of 20 SNPs with 5 SNPs of overleap in Andean populations.** Genes related to SNPs inside the 95.95th percentile of PBSn values and thye 95.95th percentile of windows PBSn mean (red line) that also present high values for XPEHH are labeled.

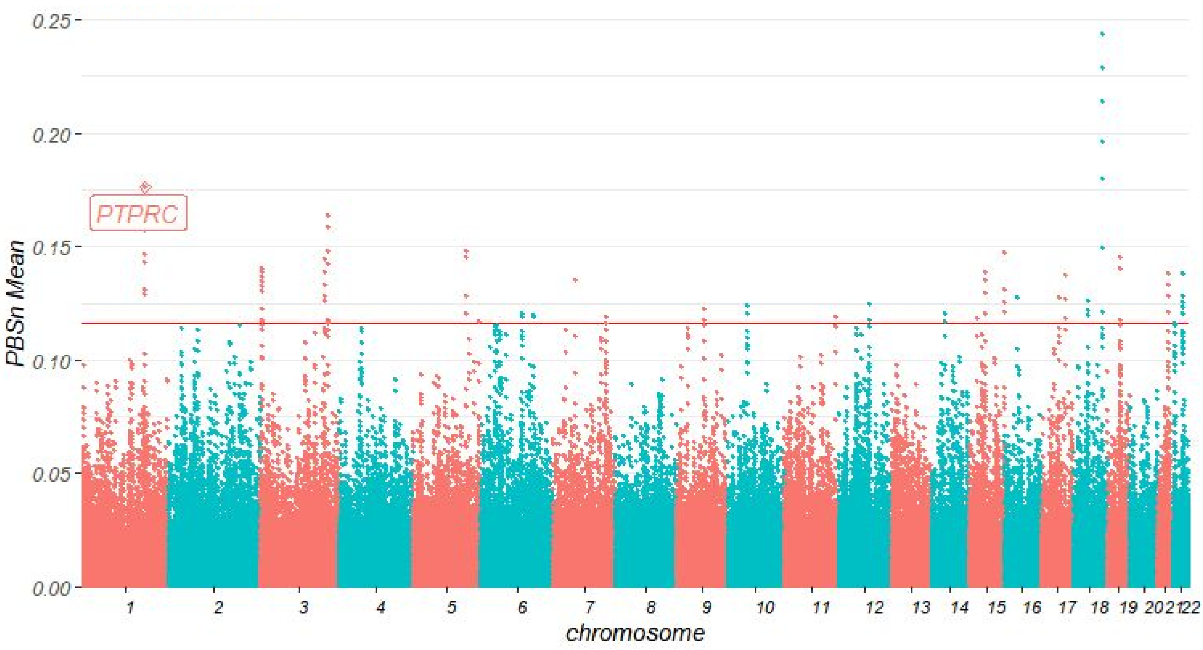


**Figure S27: PBSn mean values for windows of 20 SNPs with 5 SNPs of overleap in Amazon populations.** Genes related to SNPs inside the 95.95th percentile of PBSn values and thye 95.95th percentile of windows PBSn mean (red line) that also present high values for XPEHH are labeled.

**Supplementary Tables (In attached compressed file)**

**Table S1:** Description of 19 studied Native American populations from Peruvian National Institute of Health and from Laboratory of Human Genetic Diversity. Ashaninka population was sampled twice independently, for this reason, we merge these samples in a unique Ashaninka group and a total of 18 studied populations.

**Table S2:** List of all samples included in the Native 500K dataset.

**Table S3:** List of all samples included in the Native 230K dataset.

**Table S4:** SNPs under selection in Andean populations according to Population Branch Statistic (PBS) test.

**Table S5:** SNPs under selection in Amazon populations according to Population Branch Statistic (PBS) test.

**Table S6:** SNPs under selection in Andean populations according to Population Branch Statistic (PBS) and Cross-Population Extended Haplotype Homozygosity (XP-EHH) tests

**Table S7:** SNPs under selection in Amazon populations according to Population Branch Statistic (PBS) and Cross-Population Extended Haplotype Homozygosity (XP-EHH) tests
